# Conserved T cell receptor usage underpins recognition of CD1c presenting a mycobacterial lipid

**DOI:** 10.64898/2026.06.17.733038

**Authors:** Thinh-Phat Cao, Caroline Soliman, Samuel J. Redmond, Tori Tappen, Jade Kollmorgen, Qiji Geng, D. Branch Moody, Thomas J. Scriba, Adam P. Uldrich, Adriaan J. Minnaard, Chetan Seshadri, Hariprasad Venugopal, Adam Shahine, Jamie Rossjohn, Dale I. Godfrey, Nicholas A. Gherardin

## Abstract

The mechanism by which T cell receptors (TCR) recognise mycobacterial lipids presented by CD1 family members not well understood. We used CD1c tetramers loaded with the mycobacterial phosphomycoketide (PM) or mannosyl-PM (MPM) to isolate T cells *ex vivo* in healthy blood donors and in individuals from a tuberculosis (TB)-endemic region of South Africa, with higher frequencies observed in individuals from the TB-endemic region. High throughput analysis of >200 paired CD1c-mycoketide tetramer^+^ αβTCRs identified a conserved TCR motif, encoded by *TRBV4-1* or *TRBV7-9* variable region genes in greater than half of TCRs sequenced. A cryo-EM structure of a TRBV7-9^+^ TCR in complex with CD1c-PM demonstrated that the TCR bound to the F’ side of CD1c, directly contacting the phospholipid antigen and F’-portal residues. Analysis of multiple T cell clones interacting with CD1c mutants suggested that this TCR docking mode is representative of the larger TRBV7-9^+^ T cell population. Collectively, this study provides insight into mycobacterial lipid-antigen recognition by CD1c-restricted T cells.

**Summary:** Cao and Soliman et al study CD1c-restricted human T cells that recognize mycobacterial phosphomycoketide lipids, where they are shown to be more frequent in blood a TB-endemic region. The authors describe a strong TCR-β repertoire bias and use cryoEM to provide a high-resolution structure of an archetypal TCR engaging CD1c in complex with a mycobacterial lipid.

## Introduction

Members of the CD1 family of antigen-presenting molecules present lipid antigens to T cells (Van Rhijn et al., 2015). The most extensively studied CD1-T cell interaction is that between CD1d and type I NKT cells, which are involved in diverse settings of disease and homeostasis (Rossjohn et al., 2012). Unlike mice, humans also express the group I CD1 family members that includes CD1a, CD1b and CD1c isoforms. These molecules vary in their tissue expression, intracellular trafficking pathways and architecture of their antigen-binding clefts, thereby collectively capturing the cellular lipidome with non-redundant roles in immunity(Huang et al., 2023; Rossjohn et al., 2026; Van Rhijn et al., 2015). Yet there are many facets of CD1-mediated T cell immunity in humans that we do not understand, especially for CD1c, which is the least extensively studies CD1 antigen presenting molecule.

CD1a, CD1b and CD1c present lipid antigens from *Mycobacterium tuberculosis* (Mtb)(Kawashima et al., 2003). For example, CD1a presents the lipopeptide dideoxymycobactin (DDM), CD1b presents mycolates including mycolic acid (MA)(Beckman et al., 1994) and MA-derivatives such as glucose monomycolate (GMM)(Moody et al., 1997), glycerol monomycolate (GroMM)(Layre et al., 2009) and trehalose monomycolate (TMM)(Sakai et al., 2024), whereas CD1c can present the mycoketide lipids, phosphomycoketide (PM) and mannosyl-β-1-phosphomycoketide (MPM)(Godfrey et al., 2015; Van Rhijn et al., 2015). The extent to which mycobacteria-restricted T cells participate in host responses is unknown, but several studies suggest that CD1b-lipid-reactive T cells are expanded in the blood of active and latent tuberculosis patients(Gilleron et al., 2004; Layre et al., 2009). Also, CD1 and mycobacterial lipid-reactive T cells produce anti-mycobacterial effector molecules including granulysin(Stenger et al., 1998), TNF and IFN-γ(Gilleron et al., 2004).

CD1-lipid complexes are detected by antigen-specific T cells *via* their TCRs. For example, type I NKT cells are defined by inter-donor conserved TCRs, where an invariant TCR α-chain (*TRAV10*-*TRAJ33* in humans) underpins CD1d-lipid recognition(Borg et al., 2007). However, the extent to which pathogen-specific T cells recognising other CD1 isoforms use public TCRs is not yet understood. For example, many T cells recognising microbial lipid antigens express diverse TCRs(Van Rhijn and Moody, 2015), but two patterns for CD1b and CD1c have been described. Namely, germline-encoded mycolyl lipid-reactive (GEM) T cells encoded by *TRAV1-2-TRAJ9* TCR-α chains recognise CD1b-GMM with high affinity(Van Rhijn et al., 2013), whereas *TRBV4-1* and *TRAV17* encoded TCRs, known as ‘LDN5-like’ T cells, recognise CD1b-GMM or CD1b-trehalose monomycolate complexes with moderate affinity(Van Rhijn et al., 2014). In addition, CD1c-autoreactive T cells express *TRAV4-1*(Guo et al., 2018) and the *TRAV4-1* encoded residues that likely bind two related hydrophobic patches shared by CD1b and CD1c(Reinink et al., 2019; Sakai et al., 2024). Whether CD1c supports conserved TCRs beyond this *TRAV4-1* motif remains unknown.

CD1c-restricted T cells are present in healthy human blood, and CD1c-self-lipid antigen-reactive T cell responses have been implicated in anti-tumour responses(de Lalla et al., 2011; Gherardin et al., 2021b; Lepore et al., 2014). However, how CD1c-restricted TCRs distinguish between different lipid antigens is unknown(Matsunaga et al., 2004; Moody et al., 2000). The only published structural characterisations of a TCR binding a CD1c-lipid complex are that of the autoreactive αβTCR clone 3C8, isolated from human blood, and s2c TCR, isolated from a phage-display library, whereby lipid antigens were not directly contacted by the TCR(Szoke-Kovacs et al., 2025; Wun et al., 2018). How CD1c-restricted TCRs recognise foreign lipids thus remain unknown. Indeed, CD1c and TCR interactions may differ fundamentally from the modes used by other CD1 isoforms, as recent CD1c-lipid structures show that CD1c can present two lipids at once, with one protruding from the membrane distal, top surface of CD1c and the other extending sideways through the CD1c G’-portal (Cao et al., 2025). Thus, we investigated how two established mycobacterial lipid antigens PM and MPM(Ly et al., 2013) were recognised by TCRs.

CD1c-PM and CD1c-MPM tetramer complexes have been used to study lipid-antigen recognition and processing in CD1c-restricted T cells, which is also interesting because MPM can be de-mannosylated to generate PM(Ly et al., 2013). Unlike CD1a, CD1b and CD1d tetramers, however, the *ex-vivo* application of CD1c tetramers has been hindered by false positive staining patterns caused by the interaction between CD1c and CD36 family members(Gherardin et al., 2021a). In this study, we used CD36 blockade to perform an *ex-vivo* study on CD1c-PM and MPM antigen-reactive T cells in human blood(Gherardin et al., 2021a). We identify and characterise CD1c-restricted αβ T cells in human blood and demonstrate strong TCR repertoire biases based on the *TRBV7-9* gene, which influences fine lipid-antigen specificity. We solved the cryo-electron microscopy (cryo-EM) structure of a typical TRBV7-9^+^ TCR in complex with CD1c-PM, providing structural insight into antigen-recognition and repertoire bias by the CD1c-PM-restricted αβTCR repertoire.

## Results

### Defining reagents with precise antigen specificity

To study human CD1c-restricted lipid antigen-specific T cell populations, we first validated loading of CD1c with PM and MPM (**Figure 1A**), using established CD1c-restricted TCRs with diverse specificity. These included the CD1c-autoreactive clone, 3C8(Porcelli et al., 1992), the CD1c-MPM-specific clone, CD8.1(Ly et al., 2013; Moody et al., 2000; Rosat et al., 1999), the CD1c-PM-specific clone, DN6(Beckman et al., 1996; Ly et al., 2013; Moody et al., 2000) and an MPM and PM cross-reactive clone, 22.5(Roy et al., 2014) (**Figure 1B**). These TCRs were transiently transfected, along with CD3 subunits to induce TCR surface expression, into HEK293T cells which were subsequently stained with a panel of CD1-lipid tetramers, using SCARB1 knock out HEK293T cells to reduce SCARB1-mediated binding(Gherardin et al., 2021a) (**Figure 1C & Figure S1**). As expected, no TCRs bound CD1d-α-galactosylceramide (αGalCer) tetramers except the control type I NKT TCR clone, while the CD1c-restricted TCRs gave the expected patterns of staining based on the antigen used to load CD1c. Namely 3C8 bound weakly to CD1c that was not treated with lipid antigen and so carried only self mammalian lipids endogenous to the expression system (CD1c-endo), and other TCRs that mediated specific reactions with CD1c CD1c-PM and CD1c-MPM. Clone 22.5 bound very weakly to CD1c-endo but strongly to both CD1c-PM or CD1c-MPM. CD8.1 exhibited weak binding and only to CD1c-MPM, whereas DN6 bound to CD1c-PM with weaker binding to CD1c-MPM. Thus, the CD1c-lipid tetramers were effectively loaded with antigen and could distinguish among CD1c-lipid-specific TCR clones.

**Figure 1.**
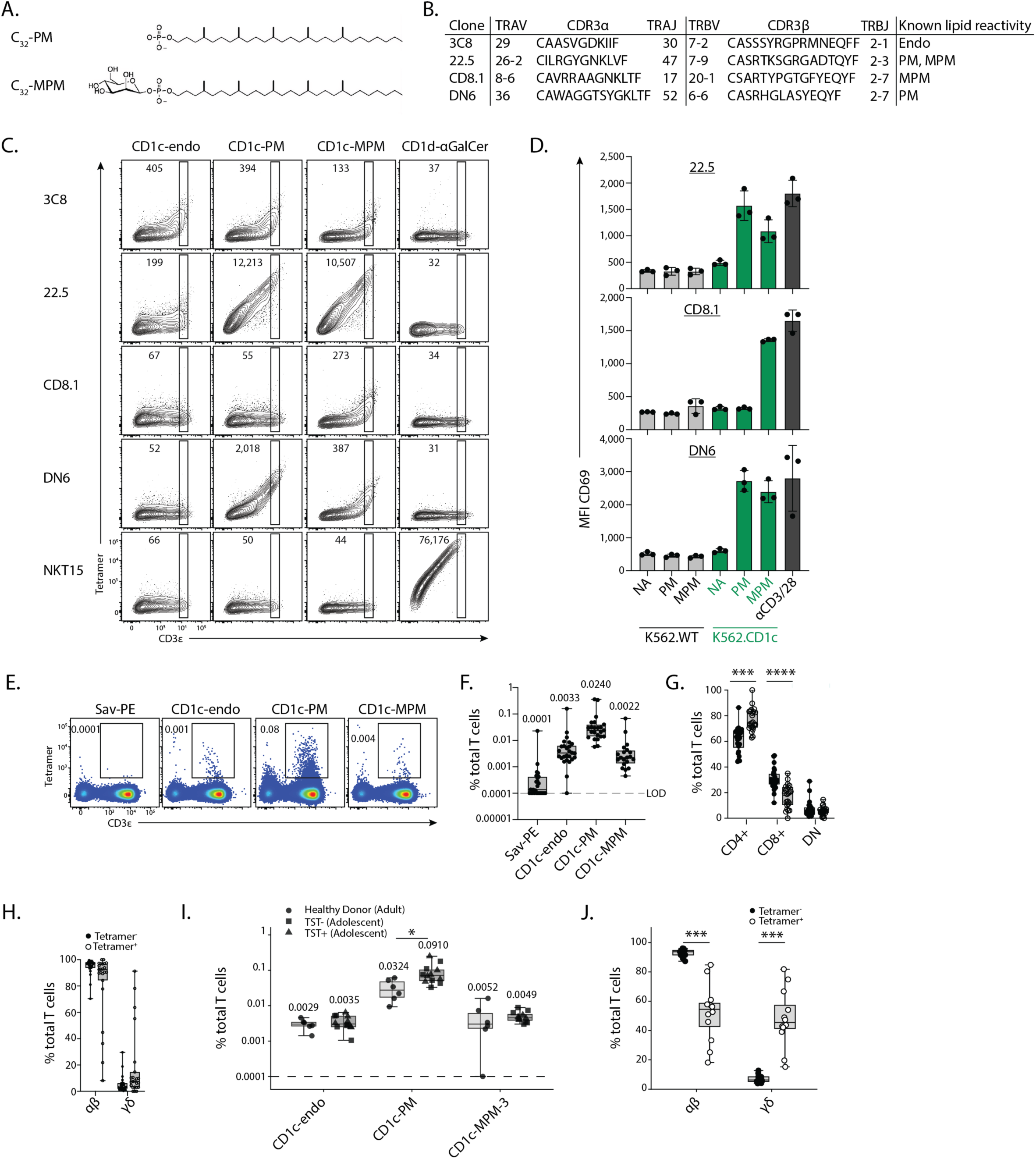
Isolation of CD1c-PM and CD1c-MPM-restricted T cells using CD1c tetramers. **A.** Chemical structures of C32-PM and C32-MPM. **B.** Table showing TCR sequences of historically isolated CD1c-restricted T cell clones. **C.** Flow cytometric contour plots showing CD1 tetramer staining on 293T.SCARB1^-/-^ cells transiently transfected to express CD1-restricted TCRs. Numbers show MFI of gated population which is consistently placed within a TCR-transfected cell line. **D.** Bar graphs showing CD69 expression on SKW-3.TCR cell lines after co-culture with WT or CD1c-overexpressing K562 cells with or without PM or MPM. **E.** Representative flow cytometric pseudo-colour plots gating on CD19^-^CD14^-^ lymphocytes after dead cell and doublet exclusion showing CD1c tetramer staining on CD3^+^ T cells. Numbers in plots depict proportion. **F-H.** Box and whisker plots showing: **F.** the proportion of T cells in healthy donor PBMC samples that stain with CD1c tetramers, **G.** the coreceptor distribution of CD1c-PM tetramer^+^ versus tetramer^-^ T cells (p<0.0001 and p<0.00001 by paired t-test), and, **H.** The proportion of αβ or γδ T cells within CD1c-PM tetramer^+^ versus tetramer^-^ T cells. **I.** Box and whisker plots showing the proportion of T cells in healthy donor PBMC samples and TST^+^ or TST^-^ South African adolescent PBMC samples that stain with CD1c tetramers. (p=0.01 by Welch’s unpaired t-test). **J.** The proportion of αβ or γδ T cells within CD1c-PM tetramer^+^ versus tetramer^-^ T cells of South African adolescent PBMC samples (combined TST^+^/TST^-^; p<0.0001 by paired t-test). Data in C is representative of n=3 independent experiments. *See also Figure S1.* Data points in D represent mean of two technical replicates from 3 independent experiments. Data F-H are derived from 26 healthy donor PBMC samples. Data in I-J are derived from 12 TST^+/-^ samples and 6 healthy donor PBMC samples. All bar graphs depict mean and SEM. Box and Whisker plots depict median, IQR and minimum and maximum points. Numbers above graphs in F and I represent mean.

To further validate these TCR specificities for antigen, stable SKW-3 cell lines were generated expressing each TCR and these were co-cultured with K562 antigen-presenting cells transduced to express CD1c compared to untransduced controls, with or without PM or MPM (**Figure 1D**). Each cell line responded to positive-control beads coated with anti-CD3 and anti-CD28 by upregulating CD69. In antigen-presentation conditions, each clone was only activated in the presence of CD1c-expressing K562 cells, demonstrating the CD1c-restriction by these TCRs. In line with tetramer staining, 22.5 and DN6 both responded to PM and MPM, noting that MPM can be converted intracellularly to PM through demannosylation(Ly et al., 2013). Clone CD8.1, on the other hand, only responded to MPM. These data validated the utility of CD1c antigen-loaded tetramers for identifying TCRs with specificity for CD1c plus particular antigens.

### Isolation of CD1c and foreign phospholipid antigen specific T cells in humans

CD1c tetramers can successfully identify T cells that are CD1c autoreactive(Wun et al., 2018), CD1c and PM-reactive or CD1c and MPM-reactive(Ly et al., 2013), but these tetramers can also stain CD36 proteins on T cells and non-T cells(Gherardin et al., 2021a). Therefore, we included anti-CD36 blockade when using CD1c tetramer staining to assess the frequency of CD1c-restricted T cells in PBMC *ex-vivo* (**Figure 1E**). As previously reported(Gherardin et al., 2021a), CD1c-endo staining was rare amongst PBMC from a cohort of healthy blood samples, although was nonetheless above the background staining with streptavidin-PE (Sav-PE) (**Figure 1F**). CD1c-MPM tetramers gave a similar staining profile and frequency as the CD1c-endo tetramer-stained T cells, whereas CD1c-PM tetramers consistently labelled more T cells, (0.02 % mean), which is a frequency that is similar to that of type I NKT cells in the periphery(Gherardin et al., 2018). However, in each case, the staining profile was a spread of cells ranging from low to high intensity (**Figure 1E**) rather than a (Gherardin et al., 2021a) population of cells similar to the clonal staining patterns associated with semi-invariant populations like type I NKT cells and mucosal-associated invariant T (MAIT) cells(Gherardin et al., 2021a). This pattern suggests detection of a heterogeneous population of T cells. Focussing on the more abundant CD1c-PM-staining cells, further analysis revealed that CD1c tetramer^+^ cells were enriched for CD4^+^ T cells relative to the CD1c tetramer^-^ population (**Figure 1G**). Furthermore, in some donors, there was a high frequency of T cells utilising γδ TCRs (**Figure 1H**). These characteristics are in line with previous studies of CD1c-restricted T cells(Gherardin et al., 2021a; Ly et al., 2013), supporting the contention that the CD1c-PM tetramer^+^ population is enriched for CD1c-restricted T cells.

### CD1c-PM-reactive T cells analysis in mycobacteria-exposed individuals

To further assess CD1c-PM-reactive T cells in a population with relevant mycobacterial antigen exposure, we stained PBMCs from a cohort of healthy adolescents residing in a high tuberculosis-burden region of South Africa (**Supplementary table S1**)(Mahomed et al., 2011). All participants received *Mycobacterium bovis* Bacille Calmette–Guérin (BCG) vaccination at birth, in accordance with South African guidelines. Participants were stratified as either tuberculin skin test and QuantiFERON®-TB Gold In-Tube positive (QFTG^+^TST^+^, hereafter TST^+^) or negative (QFTG^−^TST^−^, hereafter TST^−^), with n=6 per group. TST^+^ individuals were considered to have evidence of prior Mtb antigen exposure and presumed Mtb infection. PBMCs were stained directly *ex vivo* with CD1c-endo, CD1c-PM, and CD1c-MPM-3 tetramers, incorporating CD36 blockade to reduce nonspecific staining. CD1c-MPM-3 contains a hydrolysis-resistant MPM-3 lipid that cannot be processed to PM, enabling discrimination of PM-reactive populations from cells reactive to intact MPM(Reijneveld et al., 2021).

We compared tetramer staining in this South African adolescent cohort with a healthy adult PBMC cohort from Seattle. The Seattle adult PBMC samples showed frequencies of CD1c tetramer^+^ cells that were similar to those observed in the Australian PBMC cohort (**Figure 1I and 1F**). Within the South African adolescent cohort, TST^−^ and TST^+^ individuals displayed similar frequencies of CD1c tetramer^+^ cells and were therefore combined for subsequent analysis. Compared with Seattle PBMC samples, South African adolescent PBMCs exhibited an increased frequency of CD1c-PM tetramer^+^ cells, but not CD1c-endo or CD1c-MPM-3 tetramer^+^ cells (p=0.01 by Welch’s unpaired t-test; **Figure 1I**). This pattern is consistent with published results for CD1b-GMM-reactive T cells, where increased frequencies correlate with mycobacterial exposure rather than TB disease status(Layton et al., 2018; Lopez et al., 2020). Further analysis of CD1c-PM tetramer^+^ cells from the South African adolescent cohort revealed a mixed αβ and γδ TCR composition, with each population comprising approximately half of the tetramer^+^ cells (**Figure 1J**). These data suggest that CD1c-PM-reactive T cells may be expanded among individuals residing in TB-endemic regions.

### Antigen-specificity and TCR repertoire of CD1c-PM-restricted T cells

To further test the specificity of the CD1c-PM tetramers, and to generate more cells to study, we next used magnetic enrichment of cells from healthy donor blood-derived buffy coats using CD1c-PM tetramers (**Figure 2A**). After enrichment, the frequency of CD1c tetramer^+^ T cells was increased, and the CD3ε versus tetramer staining pattern was more compact, providing clear tetramer^+^ *versus* tetramer^-^ populations from which to examine other phenotypic markers (**Figure 2A**). Comparing co-receptor distribution on CD1c-PM-restricted T cells before and after enrichment resulted in a stronger CD4^+^ and DN T cell bias post-enrichment, providing further confidence of the isolation of CD1c-restricted T cells (**Figure 2B**). The ratio of αβ to γδ T cells was also assessed pre and post enrichment, although these did not appreciably change (**Figure 2C**).

**Figure 2.**
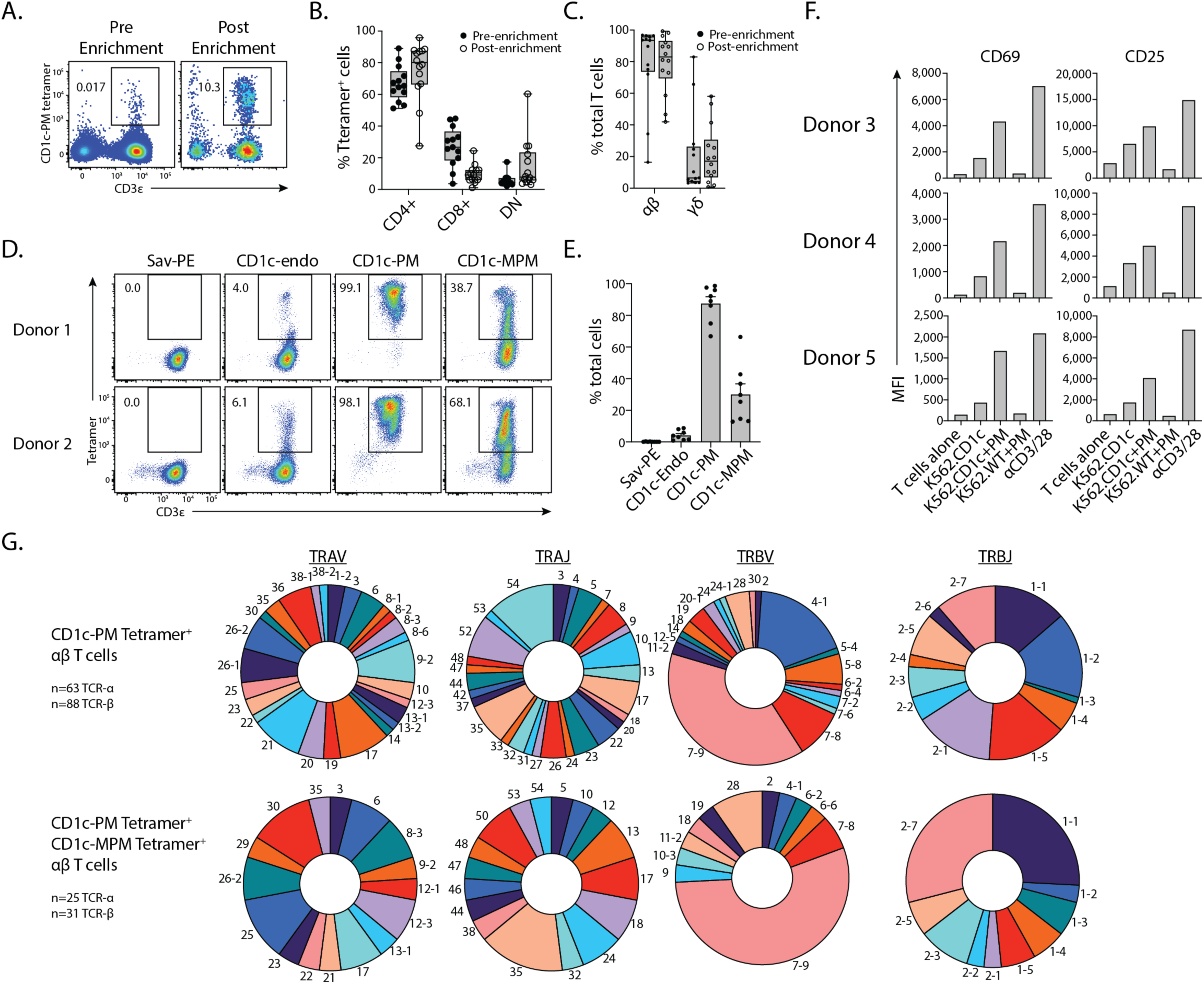
Isolation and analysis of CD1c-PM-restricted T cells. **A.** Representative flow cytometric pseudo-colour plots gating on CD19^-^CD14^-^ lymphocytes after dead cell and doublet exclusion showing CD1c-PM tetramer staining on CD3^+^ T cells before and after magnetic enrichment. **B-C.** Box and whisker plots showing: **B.** the coreceptor distribution of CD1c-PM tetramer^+^ T cells, and **C.** the proportion of αβ or γδ T cells within CD1c-PM tetramer^+^ T cells in 14 enriched versus 13 unenriched samples. **D.** Representative flow cytometric pseudo-colour plots gating on CD19^-^CD14^-^lymphocytes after dead cell and doublet exclusion showing CD1c-PM tetramer staining on CD1c-PM sort-enriched cells after *in vitro* expansion. **E.** Bar graph showing cumulative data as per D from 8 enriched and expanded samples. **F.** Bar graphs showing MFI CD69 and CD25 on polyclonal primary human CD1c-PM-restricted T cells from 3 unrelated healthy blood donors after 20 hrs co-culture with CD1c-overexpressing K562 cells with and without exogenous PM or MPM. **G**. Pie charts showing TCR gene usage of CD1c-PM sort-enriched and in vitro expanded: Cells in the top panel were single-cell sorted prior to sequencing using CD1c-PM tetramers while those in the bottom panel was sorted using CD1c-MPM tetramers and had been demonstrated to stain with CD1c-PM. CD1c-PM αβTCRs derived from 101 unique TCRs from 8 blood donors. CD1c-MPM αβTCRs derived from 33 unique TCRs from 4 blood donors. *See also Tables S2 and S3.* All bar graphs depict mean, and SEM is shown in C and E. Box and Whisker plots depict median, IQR and minimum and maximum points. Numbers in flow cytometry plots depict proportion

In another test to validate the specificity of CD1c-PM tetramer^+^ cells, magnetically-enriched cells were FACS-sorted, expanded *in vitro* using anti-CD3 and anti-CD28 antibodies over 1-2 weeks, and subsequently re-stained with distinct tetramers (**Figure 2D-E**). In most donors, most cells, 90% on average, re-stained with CD1c-PM tetramers, highlighting the reliability of these tetramers to capture CD1c-PM-reactive T cells. For example, most of these cells failed to stain with CD1c-endo tetramers demonstrating PM antigen-specificity and showing that these CD1c tetramers are not isolating CD1c-autoreactive T cells. A lower proportion of these cells, however, also bound CD1c-MPM tetramers, typically with lower staining intensity, suggesting some cross-reactivity within the CD1c-PM-restricted T cell repertoire.

Finally, to assess if these polyclonal pools of CD1c-PM tetramer^+^ cells could indeed respond to CD1c-PM antigen-presentation, they were co-cultured with wildtype or CD1c-expressing K562 cells, with or without PM, and assessed for CD69 and CD25 upregulation (**Figure 2F**). This analysis was limited to 3 donors, as T cells from the remaining donors were confounded by high basal CD69 and CD25 expression post *in vitro* expansion. Nonetheless, in all three donors assessed, each pool responded to positive control beads coated with anti-CD3 and anti-CD28 whereas neither responded to wildtype K562 cells. CD1c-expressing K562 cells induced some reactivity in each pool, however, all three pools exhibited enhanced CD69 and CD25 in response to exogenous PM, further confirming PM-dependent recognition by these cells.

### CD1c-restricted TCR motifs

To quantitatively assess the TCR-repertoire of CD1c-PM-restricted αβ T cells, we sorted single cells from polyclonal pools of *in vitro* expanded αβ T cells using either CD1c-PM or CD1c-MPM tetramers. Two hundred and seven paired TCR-α and TCR-β chains were obtained after sequencing (**Figure 2G, Supplementary tables 2-3**). CD1c-PM tetramer-sorted αβ T cells (**Figure 2G**) had highly variable TCR-α chains with no clear bias in *TRAV* or *TRAJ* gene usage, nor CDR3α length or motif (**Supplementary table 2**). There was a clear bias, however, in the *TRBV* usage (**Figure 2G**), with *TRBV4-1* and *TRBV7-9*, the latter gene not previously identified as enriched in CD1c-autoreactive T cells, collectively representing more than 50% of unique sequences, although CDR3β sequences were highly variable even within the *TRBV4-1* and *TRBV7-9* subgroups (**Supplementary table 2**). CD1c-MPM tetramer-sorted αβ T cells, which also stained for CD1c-PM tetramer, exhibited similar TCR-α chain diversity, however the *TRBV7-9* bias was stronger and there was no notable bias toward *TRBV4-1* unlike the CD1c-PM tetramer-sorted population. Moreover, more than half of the CD1c-MPM tetramer-sorted clones utilised *TRBJ2-7* or *TRBJ1-1* which represented a greater enrichment relative to the CD1c-PM tetramer-sorted cells (**Figure 2G and supplementary table 3**).

Accordingly, these data show that diversity is permitted within the CD1c-PM-restricted T cell repertoire, however, there is also bias toward *TRBV4-1* as has been previously seen for CD1c-autoreactive cells(Guo et al., 2018). Furthermore, we identified strong bias toward usage of the *TRBV7-9* segment with 54 of 119 unique TCRβ chains sequenced encoded this TRBV gene segment (**Supplementary table 2**). Although several TRBV7-9^+^ clones were known from a small analysis of individual T cell clones(Ly et al., 2013), this strong bias, revealed through high throughput analysis, to a single *TRBV* gene among hundreds of TCRs was notable and unexpected, warranting further study of *TRBV7-9*^+^ TCRs.

### Functional validation of the specificity of TRBV4-1 and TRBV7-9 TCR motifs

To validate the antigen-specificity of these PM and MPM-reactive αβTCRs, we selected a series of clones that were sorted with either CD1c-PM or CD1c-MPM tetramers, including three *TRBV4-1*^+^ TCRs, five *TRBV7-9*^+^ TCRs, as well as a *TRBV2^+^* TCR and *TRBV20-1^+^* TCR (**Figure 3A**). These TCRs were cloned and transiently transfected, with CD3 subunits, into HEK293T.SCARB1^-/-^ cells for surface TCR expression and then stained with a panel of CD1 tetramers (**Figure 3B**). In parallel, TCR transfectants for an NKT TCR (NKT 15) and three existing TCR transfectants with known CD1c and PM reactivity, clones 22.5, CD8.1 and DN6, were included as controls (**Figure S2**). As expected, only the CD1d-restricted control NKT15 TCR stained with CD1d-αGalCer tetramer (**Figure 3B, Figure S3**). All *TRBV4-1*^+^ and *TRBV7-9*^+^ TCRs (clones PM4.1, PM4.2, PM7.1, PM7.2, PM7.3, PM7.4) stained with CD1c-PM tetramers. However, the *TRBV2* and *TRBV20-1* (clones PM2, PM20) failed to stain with any tetramers suggesting that they may be of very low affinity or false positives from the sorting process. At best, only very weak staining was observed in some clones with CD1c-endo and vehicle tetramers, confirming the specificity for CD1c-PM that was observed in the primary cells (**Figure 2D**). Many clones showed cross-reactivity for PM and MPM, although in each case staining was brighter for PM than MPM.

**Figure 3.**
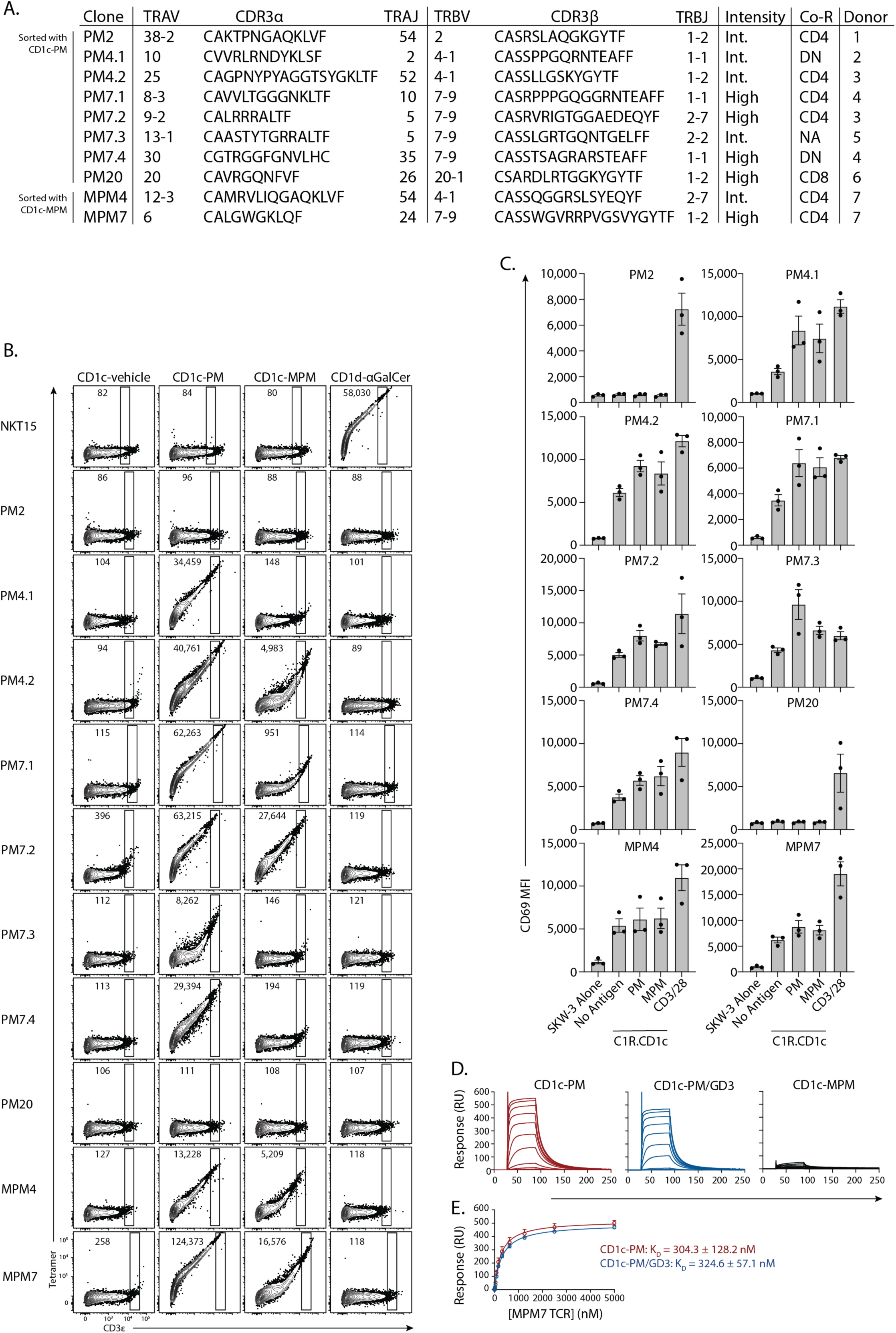
CD1c-antigen-reactivity by αβ T cells. **A.** Table showing TCR sequences of cloned CD1c-PM-restricted αβ T cell clones. **B.** Flow cytometric contour plots showing CD1 tetramer staining on HEK293T.SCARB1^-/-^ cells transiently transfected to express CD1-restricted TCRs. Numbers in plots depict proportion. **C.** Bar graphs showing CD69 expression on SKW-3.TCR cell lines after co-culture with CD1c-overexpressing C1R cells. Data in B is representative of n=2 independent experiments with bars depicting mean and SEM. Data points in C represent mean of two technical replicates from 3 independent experiments. *See also Figures S2, S3 and S4. **D.*** Binding analysis of MPM7 TCR to CD1c-PM, CD1c-PM-GD3, and CD1c-MPM using surface plasmon resonance. Representative steady-state sensorgrams of the three binding analyses are shown. K_D_ values (mean ± S.D.) in the equilibrium plot were obtained from n = 4 independent experiments.

We next asked whether any of these CD1c-PM-reactive TCRs could cross-react with other mammalian lipids. To assess this, we stained the panel of TCR-transfected HEK293T.SCARB1^-/-^clones with a panel of CD1c tetramers loaded with self-lipids including ganglioside GD3, sulfatide or a series of phospholipids including phosphatidic acid (PA), phosphatidylcholine (PC), phosphatidylethanolamine (PE), phosphatidylglycerol (PG), phosphatidylinositol (PI) or phosphatidylserine (PS; **Figure S4**). Most TCRs lacked detectable binding to any of these CD1c-Ag tetramers, and MPM7 showed a very weak response. In some instances, however, there were subtle modulations in staining, for example, the weak CD1c-endo tetramer staining of clones 22.5 and PM7.2 was reduced with CD1c-GD3 and CD1c-sulfatide tetramers.

To probe whether this antigen-recognition translated to a TCR-mediated signalling event, stable TCR-expressing SKW-3 (β2M^-/-^) cell lines were produced, and these were co-cultured with CD1c-transduced C1R antigen-presenting cells (**Figure 3C**). While clones PM2 and PM20 failed to react to CD1c in the presence or absence of exogenous lipid, all other TCRs demonstrated a degree of inherent CD1c autoreactivity. This likely reflects the much higher expression, and therefore avidity, of CD1c in a transduced cellular setting, allowing a degree of TCR-dependent activation even in the absence of the defined lipid antigens. Activation by all other clones, aside from clone MPM4, was increased in the presence of exogenous PM and MPM to varying degrees, though this was not as pronounced as was observed with the primary cells (**Figure 2F**). This data collectively validates the antigen-specificity of the TCRs and their ability to signal in response to CD1c and antigen.

### Biochemical characterisation of αβTCR binding to CD1c-PM

To date, formal structural analysis of αβTCRs binding to CD1c-mycobacterial lipid antigen complexes has not been reported. We focused on the MPM7 TCR as it utilised the unexpectedly enriched *TRBV7-9* TCR-β chain and exhibited the highest tetramer staining intensity (**Figure 3B**). Binding analysis using surface plasmon resonance (SPR) demonstrated an affinity of MPM7 for CD1c-PM of K_D_ = 304.3 ± 128.2 nM (**Figure 3D-E**). This is considered very strong relative to classical αβTCR binding of peptide-MHC(Rossjohn et al., 2015), more closely aligning to that of the high affinity interaction between the type I NKT TCR and CD1d-αGalCer(Kjer-Nielsen et al., 2006). In contrast, no binding was observed between the MPM7 TCR and CD1c-MPM, indicating that the TCR is specific for the non-mannosylated mycoketide, despite the observation of binding when using CD1c-MPM tetramers (**Figure 3B**). This is likely due to a fraction of the MPM molecules degrading to PM through phosphate ester hydrolysis to enable low intensity binding when using tetramers(Reijneveld et al., 2021). This is furthermore in line with the weak CD1c-MPM tetramer staining of DN6 (**Figure 1B**), which is reported to be PM specific(Ly et al., 2013), and the relatively weak CD1c-MPM tetramer staining on the CD8.1 TCR (**Figure 1B**).

We recently demonstrated that lipids with large bulky headgroups such as gangliosides are displayed in a sideways fashion, with the bulky headgroup protruding laterally through a side G’-portal of CD1c, and that this sideways presentation was found to be compatible with co-presentation of a second lipid via the apical portals, including PM(Cao et al., 2025). To determine whether sideways-presented lipids could alter antigen-recognition of CD1c-PM-specific TCRs, SPR were also performed using CD1c co-loaded with both PM and GD3. Here, the binding affinity was essentially unchanged relative to PM only (324.6 ± 57.1 nM, **Figure 3D-E**). Thus, the presumably sideways-oriented GD3 does not affect MPM7 TCR recognition of CD1c-PM.

### Structural overview of αβ TCR recognition of CD1c-PM

We next determined the 2.9 Å resolution structure of a *TRBV7-9*^+^ TCR in complex with CD1c-PM and GD3 using cryo-electron microscopy (cryo-EM) (**Figure S5 and Supplementary Table 4**). The MPM7 TCR docked on the membrane distal surface known as the A’-roof that represents the’top’ rather than the side of CD1c. This surface, which covers the top of the A’-pocket, provides a substantial surface for TCR contacts with CD1c that do not directly involve the bound PM ligand (**Figure 4 and Figure S6A**). A clear well-defined density corresponding to PM occupied the A’-pocket, located under the A’-roof, with the local resolution approaching ∼ 2.6 Å (**Figure S6B**).

**Figure 4.**
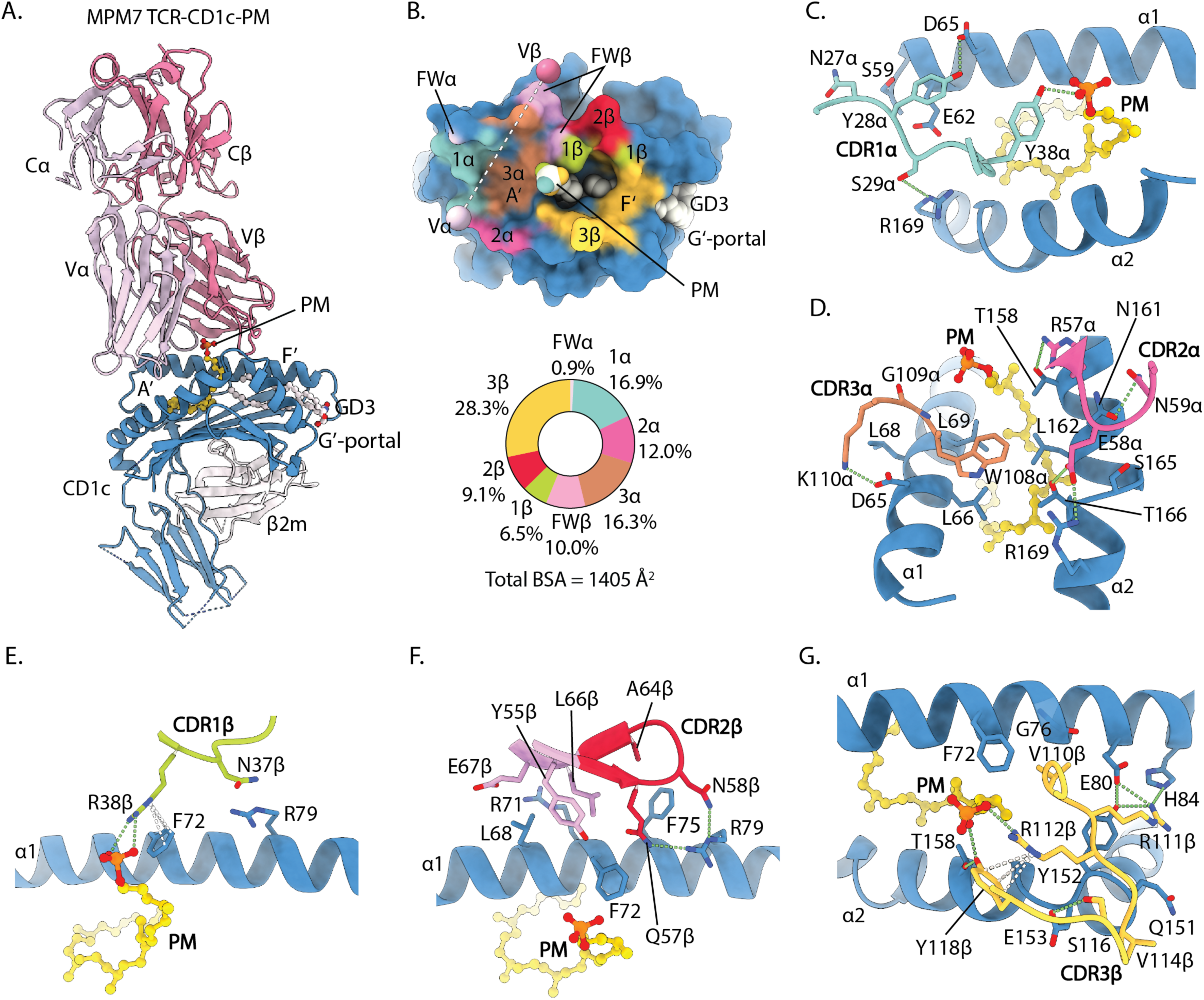
Characterization of the MPM7 TCR-CD1c-PM interaction. **A.** Overall docking topology of MPM7 TCR atop CD1c-PM. **B.** Footprints of MPM7 TCRs on the surface of CD1c, and pie-charts demonstrating the relative contributions of TCR regions to the total buried surface area (BSA) with CD1c-PM in the complex. **C.** Details of interactions of CDR1α to CD1c-PM. CDR1α was positioned along the midline of the antigen-binding cleft, where S29α and Y28α mediated hydrogen bonds with R169 and D65, respectively, while N27α and Y28α formed van der Waals (vdW) contacts with S59 and E62 of CD1c **D.** Details of interactions of CDR2α and CDR3α to CD1c-PM. CDR2α engaged CD1c via three hydrogen bonds between R57α-T158, E58α-R169, and N59α-N161, with E58α and N59α further forming vdW interactions with S165 of the α2-helix of CD1c While CDR3α engaged CD1c via a single salt bridge between K110α-D65, a vdW interaction between G109α-L68, and hydrophobic interactions between W108a and a cluster comprising L66, L69, L162, and T166 from CD1c. Notably, K110α is a TRAJ-gene encoded lysine that commonly occurs in the 3’ sequence of *TRAJ* genes. **E.** Details of interactions of CDR1β to CD1c-PM. **F.** Details of interactions of CDR2β (red) and the FWβ (pink) regions to CD1c. **G.** Details of interactions of CDR3β to CD1c-PM. Hydrogen bonds and π-interactions are depicted as green and white dashed-lines, respectively.

The other internal pocket, known as the F’-pocket, contained a separate continuous lipid density consistent with the two carbon-tails of GD3 (**Figure S6B**). The GD3 glycan headgroup, which was expected to protrude sideways through a lateral gap in the F’-pocket wall, known as the G’-portal(Cao et al., 2025), was not observed, likely due to the intrinsic flexibility of the glycan when located outside the constraints of the CD1c cleft (**Figure S5**). A ceramide backbone was modelled into this density, assigning a 36:1 tail length, in an orientation similar to that reported previously(Cao et al., 2025). The vertically oriented MPM7 TCR that bound on the membrane distal surface of CD1c did not contact the lipid tails of GD3 in the F’-pocket, and the TCR docking location was distant from the expected protrusion site of the GD3 headgroup. Overall, this general orientation of the TCR to CD1c and the two carried lipids identified a’vertical’ approach to the membrane distal surface of CD1c and contact with the single chain PM antigen.

The MPM7 TCR docked 62° across the long axis of CD1c-PM with a total buried surface area (BSA) of 1405 Å^2^, of which the TCR α-and β-chains contributed approximately equally (46.1 % and 53.9 %, respectively) (**Figure 4A-B**). Among the complementarity determining regions (CDR) regions, the footprint was dominated by CDR3β (28.3 %), whereas CDR1β accounted for the smallest contribution (6.5 %). The remaining CDR-mediated interface was distributed among CDR1α (16.9 %), CDR3α (16.3 %), CDR2α (12.0 %), and CDR2β (9.1 %). The framework (FW) regions contributed a total of 10.9 % of the BSA, of which FWβ took up a significant portion (10.0 %). The TCR footprint covered a broad surface surrounding the A’-roof and enveloped the protruding PM headgroup emerging from the F′-portal (**Figure 4A-D & Figure S6B**). In general, the TCR α-chain’s germline-encoded regions primarily engaged the rim of the A’-roof of CD1c. (**Figure 4B-C**).

### TRBV7-9^+^ TCR β-chain-CD1c interactions

The germline-encoded elements of the TRBV7-9^+^ β-chain were bound on the α1-helix and located near to the point of antigen protrusion from the F’-pocket, which is known as the F’-portal of CD1c (**Figure 4B, E-F**). CDR1β formed van der Waals (vdW) interactionw between N37β-R79 on the α1-helix of CD1c (**Figure 4E**), and R79 was further hydrogen-bonded to the germline-encoded CDR2β region via two residues, Q57β and N58β (**Figure 4F**). F72 and F75 of CD1c also formed a concave hydrophobic cluster that mediated interactions with Q57β and A64β of the TRBV7-9^+^ β-chain (**Figure 4F & Figure S6A**). Notably, three other residues in the FWβ regions of the TCR β-chain were observed to contact CD1c, whereby Y55β and L66β formed a hydrophobic cluster with L68, R71, and F72 of CD1c, while E67β FWβ made vdW interactions with L68 and R71 of CD1c (**Figure 4F** and **S7A**).

The non-germline-encoded CDR3β loop primarily interacted in the vicinity of the F’-portal (**Figure 4B, G**), which surrounds the protruding antigen headgroup. The CDR3β loop mediated hydrophobic interactions involving V110β and V114β from CDR3β interacting with F72, G76 and the main chains of Q151 and Y152 from CD1c. In addition, hydrogen bonds were formed between S116β-E153 and R111β with the cluster comprising E80, H84, and Y152 from CD1c (**Figure 4G**). Notably, the interaction between CDR3β R111β and E80, together with the proximal Y152 and H84 on CD1c, reflects a conserved interaction also seen in CD1b-TCR complexes(Shahine et al., 2017) (**Figure S7B**). Accordingly, we provide a structural basis for the TRBV7-9^+^ β-chain bias underpinning the CD1c-restricted response, where the TCR binds on the membrane distal surface of CD1c near the centre of the platform and contacts both the antigen and F’-portal regions.

### TCR specificity toward PM

Upon MPM7 TCR ligation, CD1c underwent a marked conformational rearrangement, evident from alignment of the ternary MPM7 TCR-CD1c-PM complex with the binary CD1c-PM structure(Cao et al., 2025) (**Figure 5A**). This significant realignment is notable because CD1c is known to have conformational flexibility(Cao et al., 2025; Scharf et al., 2010) and such large changes have not generally been seen in apo versus liganded structures of other CD1 isoforms. Namely, both the α1-and α2-helices of CD1c were displaced downward, shifting by 2.9 and 2.2 Å, respectively (**Figure 5A**). These movements compressed the entire binding pocket of CD1c from 2170 to 1700 Å^3^. Consequently, the PM molecule shifted upward by 5 Å, becoming fully exposed for engagement by the TCR’s ‘cationic cup’, which is a small positively charged pocket formed by a tetrad of residues, Y38α (CDR1α), R38β (CDR1β), R112β and Y118β (CDR3β) of the MPM7 TCR (**Figure 5B, C**). Notably, R38β and R112β from the TCR adopted a further stabilised orientation through π-cation interactions with F72 of CD1c and Y118β from the TCR, respectively (**Figure 4E, G** and **5C**). A similar ‘cationic cup’ architecture has previously been observed in TCR-CD1b-lipid interactions with anionic phospholipids(Farquhar et al., 2023; Shahine et al., 2019; Shahine et al., 2017). In contrast, in autoreactive TCR-CD1c interactions, including the TRBV7-2^+^ 3C8 and TRBV7-9^+^ s2c TCRs, this ‘cationic cup’ was not formed owing to the absence of a requirement for binding a negatively charged lipid headgroup(Szoke-Kovacs et al., 2025) (**Figure S7C**). Together with a constellation of other surrounding residues, the ‘cationic cup’ configuration in the MPM7 TCR is incompatible with accommodation of the mannosyl headgroup of MPM (**Figure S7D**), thereby providing a clear structural basis for the PM-bias by the MPM7 TCR, and possible explanation for the broader preference of many CD1c-reactive TCRs to PM versus MPM.

**Figure 5.**
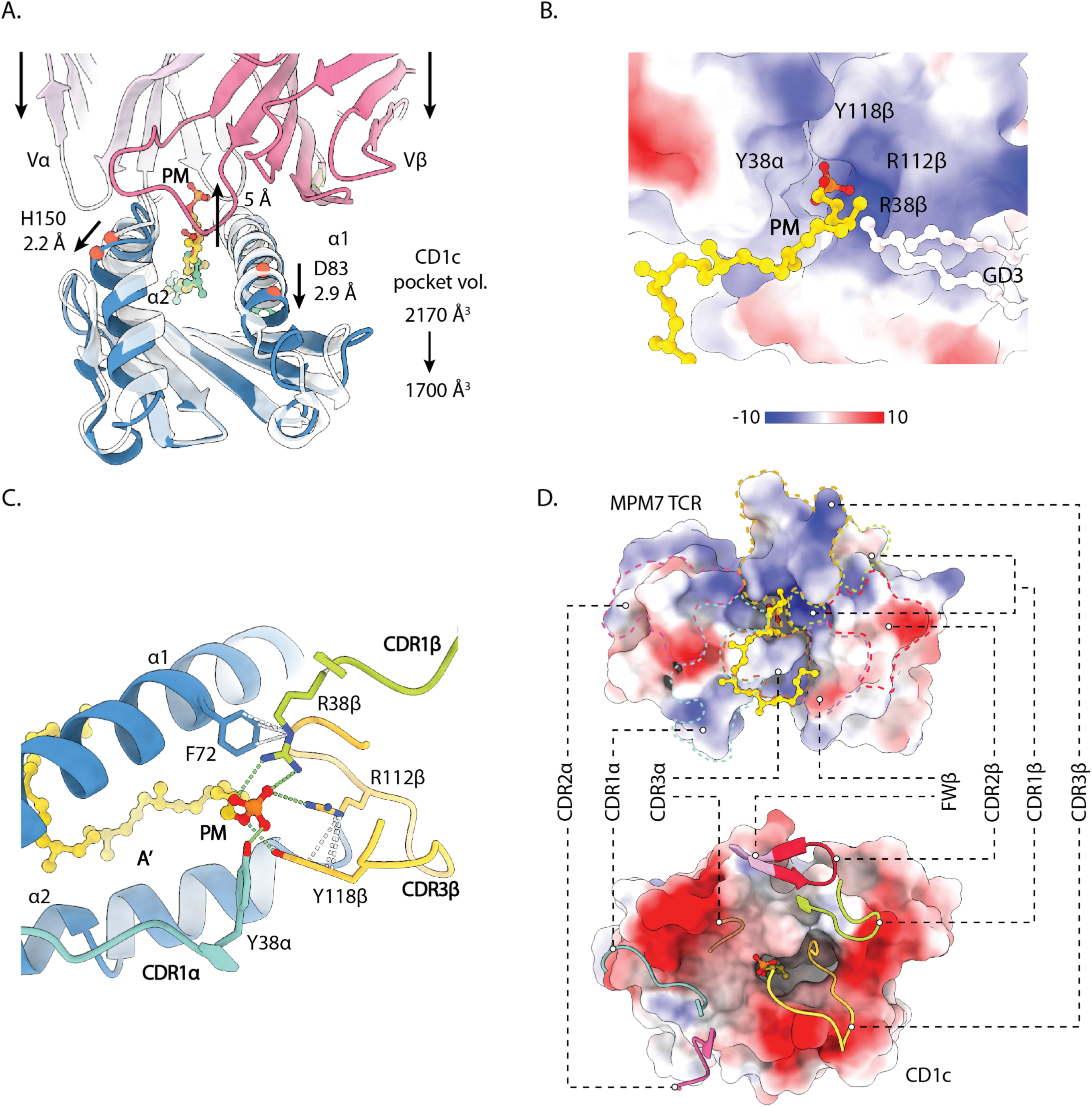
Structural basis for CD1c-PM specificity. **A.** Conformational changes of CD1c-PM upon MPM7 TCR binding. Arrows indicate direction of movement. The CD1c from the ternary structure is coloured blue, whereas the CD1c-PM binary structure (PDB ID: 9OHY) is shown in white. PM in the ternary structure and in the binary structure are depicted as yellow and green, respectively, ball-and-stick. Movements of the α1 and α2 helices were measured at the D83 Cα and H150 Cα, respectively. Movement of PM was measured at the phosphate P atom. Pocket volumes with and without MPM7 TCR engagement were calculated using CASTpFold(Ye et al., 2024). **B.** Electrostatic potential distribution on the MPM7 TCR was visualised by ChimeraX, revealing the ‘cationic cup’ that encapsulates the electronegative phosphate headgroup of the PM. **C.** Details of interaction between the MPM7 TCR to the PM headgroup. H-bonds are depicted as green dashed-lines, whereas π-cation interaction are depicted as white dashed-lines. **D.** Complementary and electrostatic surface analysis at the interface of MPM7 TCR (top) versus CD1c (bottom). PM molecule (yellow ball-and-stick) is displayed in both proteins as a marker. CDR and FWβ regions are depicted as ribbons using the same colour-codes as in Figure 4, and their corresponding positions on the MPM7 TCR surface are circumscribed by consistently colour-coded dashed-lines.

Three of the PM-bound residues from the cationic cup are from the TCR β-chain, while the sole TCR α-chain contribution is the germline-encoded Y38α. Yet, this tyrosine may not be strictly required for PM specificity, as it is not conserved among other PM-reactive TCRs such as DN6, PM7.2, PM4.1, and PM4.2 (**Figure S8A**). Conversely, the *TRBV7-9* germline-encoded R38β appears a key requirement for PM specificity, as it is mostly present in the PM-reactive TCRs but absent from others (**Figure S8B**). This PM specificity is accompanied by interface features consistent with high affinity binding of MPM7 TCR and CD1c. The calculated shape complementarity statistic, Sc(Lawrence and Colman, 1993), between these two proteins is 0.71, clearly indicating a tight complementary interface regardless of the PM molecule. Moreover, the binding surfaces of these two proteins are highly electrostatically compatible, particularly at the interfaces formed by CDR3α and CDR3β, highlighting the important contribution of these hypervariable regions (**Figure 5D**). Collectively, a series of germline-encoded and non-germline-encoded residues arising predominantly from the TCR-β chain of the MPM7 TCR represent the specificity governing determinants underpinning high affinity PM recognition when presented by CD1c (Gherardin et al., 2021a), which provides insight into TCR-β sequence conservation among CD1c and phospholipid-reactive TCRs

### Alanine-scanning CD1c mutant analysis of αβTCR docking on CD1c-PM

We next sought to assess the energetic footprint of TCR docking atop CD1c by the CD1c-PM-reactive αβTCRs. We used TCR-transduced SKW-3 cells responding to exogenous PM, presented by a panel of 14 CD1c-transduced C1R cell lines in which solvent-exposed residues along the α1-and α2-helices of CD1c had been mutated to alanine(Wun et al., 2018) (**Figure 6A**). Clone MPM7, which has the strongest intensity of CD1c-PM tetramer staining, was the only TCR where no alanine mutation reduced activation by more than 50 %. All other TCR-expressing clones exhibited recognition ‘hotspots’ atop CD1c, defined as the loss of > 50 % binding upon mutation. Further, the pattern of hotspots was similar across the TCRs, where they were positioned on the central portion of the membrane distal end of CD1c or shifted laterally toward the A’ side of CD1c, near to the point of antigen protrusion. The F72A mutation was important for recognition by all clones aside from MPM7, and others were also affected by combinations of L68, R79 and Y152. For example, clones DN6 and PM4.1 were also affected by R79A and Y152A mutations; clone PM4.2 was affected by R79A; clones PM7.1, PM7.4 and 22.5 were affected by L68A; and clones PM7.2 and PM7.3 were affected by Y152A. Thus, diverse CD1c-PM-restricted TCRs utilise similar, antigen-adjacent docking footprints to facilitate antigen recognition, and they all differ from the known hotspots of CD1c-autoreactive TCRs on the unliganded surface of CD1c(Wun et al., 2018).

**Figure 6.**
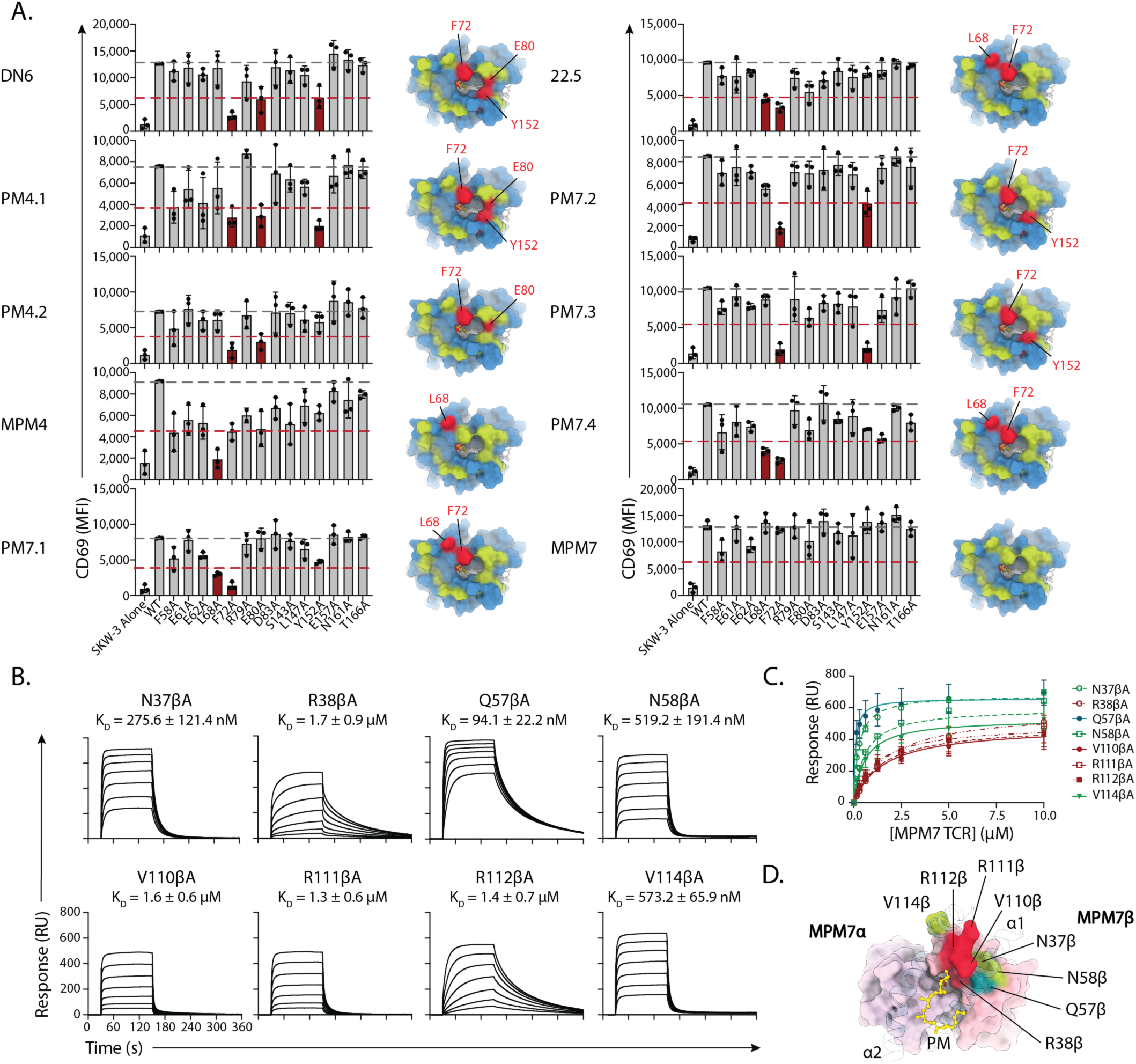
Alanine-scanning mutagenesis of CD1c-restricted TCR docking atop CD1c. **A.** Left: Bar graphs showing CD69 expression on SKW-3.TCR cell lines after 20 hr coculture with PM lipid and C1R cell lines expressing CD1c WT or alanine point mutants. Data points represent mean of two technical replicates from 3 independent experiments with bar graphs showing mean and SEM. Right: Surface representations of the top-down view of the CD1c-PM antigen-binding cleft highlighting the location of mutations that resulted in 50% or more reduction (red patches) in CD69 expression. CD1c-PM is coloured in the same colour-code with Figure 4 and 5. Green patches indicate the location of mutations that do not affect the recognition. **B,C.** Binding analysis of MPM7 TCR mutants to CD1c-PM using surface plasmon resonance. **B.** Representative steady-state sensorgrams of the 8 binding analyses are shown. K_D_ values (mean ± S.D.) were obtained from n = 2 independent experiments. **C.** Steady-state equilibrium plot of 8 binding reactions shown in B. **D.** Surface representation of MPM7 TCR showing location of the selected mutations. CD1c α1 and α2 helices are shown in transparent cartoon. Green, red, and teal footprints indicate the mutations that: do not affect; four-to five-fold reduce; and three-fold increase, the binding affinity, respectively, in consistency with panel C.

Next, soluble recombinant MPM7 TCRs were produced with alanine mutations at selected sites based on the high-resolution structure, and tested for binding to immobilised CD1c-PM by SPR. These experiments revealed that mutations at N37β and N58β from the TRBV7-9 TCR-β chain reduced binding affinity by approximately half for the MPM7 TCR-CD1c-PM interaction, whereas R38β from the TRBV7-9 TCR-β chain decreased the affinity by 4-fold (**Figure 6B**). Notably, Q57βA enhanced affinity by 4-fold, and demonstrated extended association and dissociation phases (**Figure 6C**). Mutation at V110β, R111β, and R112β on MPM7 TCR-β chain decreased the affinity with CD1c-PM by 3-fold (**Figure 6C**). Mutation at R112β also reduced the response and altered the interaction kinetics observed in the SPR sensorgram, underscoring the importance of PM headgroup contacts in TCR recognition. Interestingly, the unexpected reduction in response and affinity with CD1c-PM of the two mutants V110βA and V114βA from the CDR3β loop, despite only modest changes in hydrophobicity, suggests a stringent requirement for precise complementarity between the CDR3β loop and CD1c to achieve optimal recognition.

## Discussion

In this study, using CD1c-PM as an exemplar of a CD1 molecule presenting a foreign lipid from a medically relevant bacterium, we identified an unexpected and strongly penetrant *TRBV7-9* TCR bias, providing evidence for a public TCR in the human CD1c system. The use of public TCRs present across unrelated donors is a key feature of the type I NKT-CD1d axis, and is increasingly appreciated for the CD1b system(Reinink et al., 2019; Sakai et al., 2024; Van Rhijn et al., 2014; Van Rhijn et al., 2013). Yet, much less is known about whether similar conservation occurs within CD1c reactive TCRs. We and others have found that two TCR variable region genes are overexpressed in polyclonal T cells or multiple T cell clones that recognise CD1b or CD1c(Gherardin et al., 2021a; Guo et al., 2018; Reinink et al., 2019; Sakai et al., 2024). Specifically, *TRBV4-1* is expressed on CD1c autoreactive(Gherardin et al., 2021a; Guo et al., 2018), CD1b-GMM(Van Rhijn et al., 2014) and TMM-reactive T cells(Sakai et al., 2024), and *TRAV17* has been detected on CD1c tetramer sorted and CD1b-GMM-reactive cells(Van Rhijn et al., 2014). These emergent patterns of *TRBV* and *TRAV* genes being enriched in TCR repertoires restricted to multiple CD1 isoforms do not necessarily suggest TCR cross-reactivity to two isoforms, but they point to an emerging understanding that two monomorphic CD1 molecules may share contact mechanisms via TCRs. For example, *TRBV4-1* encoded regions of the TCR-β chain interact with a hydrophilic patch on CD1b that is it shared by CD1c(Reinink et al., 2019; Sakai et al., 2024). These gene associations however have not been validated in high throughput, and the role of lipid antigens in these interactions remained unknown.

Here, single cell TCR sequencing of polyclonal CD1c-PM-restricted T cells from healthy donors revealed multiple biases in the TCR repertoire. Firstly, we noted a frequent usage of *TRBV4-1* as has been observed amongst CD1c-endo-reactive TCRs(Gherardin et al., 2021a). That this extends to those TCRs recognising CD1c presenting a specific foreign lipid suggests that *TRBV4-1*-mediated CD1 docking may facilitate recognition of diverse lipid antigens. However, we did not observe an enrichment of *TRBV4-1* in the CD1c-PM-reactive repertoire of cells that were sorted based on CD1c-MPM staining, which may suggest that the specific docking of *TRBV4-1*^+^ TCRs on CD1c is less compatible with cross recognition of MPM. In contrast, for both PM and MPM reactive cells, we observed a strong TCR bias towards the use of *TRBV7-9^+^*a gene not observed to be enriched in the CD1c-endo-restricted T cell repertoire. A previous study that cloned five CD1c-PM-reactive TCRs by limiting dilution of CD1c-PM tetramer^+^ cells also noted that four of the five TCRs utilised TRBV7-9(Roy et al., 2014). Here, we extend that finding to a large sample size including many donors and confirm this as a clear TCR repertoire bias for these cells.

These Variable region gene biases are distinct to the semi-invariant TCR-α chains of the innate-like CD1d-αGalCer-reactive type I NKT(Borg et al., 2007; Kawano, 1997; Lantz and Bendelac, 1994) and MR1-5-OP-RU-reactive MAIT cells(Patel et al., 2013; Porcelli et al., 1993; Tilloy et al., 1999) which exhibit more restricted repertoires, pairing an almost fixed TCR-α chain with restricted *TRAV* and *TRAJ* genes and CDR3α motifs with TCR-β chains using biased *TRBV* repertoires(Godfrey et al., 2015). This is perhaps driven by a broader antigenic repertoire for the TCR-β chain biases observed with the CD1b-and CD1c-restricted populations, with the enriched germline-encoded residues driving CD1 docking and the diversity in CDR3 regions and paired variable domain fine tuning docking and allowing for lipid headgroup diversity. This conforms to the notion that TCR-repertoire bias operates on two distinct levels as has been proposed within the CD1b-GMM-reactive repertoire for the semi-invariant GEM TCRs and the LDN5-like TCRs that exhibit variable gene enrichment(Van Rhijn et al., 2014; Van Rhijn et al., 2013). How this hierarchy manifests at a functional level remains unclear. While the more restricted type I NKT and MAIT cell populations exhibit innate-like phenotypes, acquired during development(Pellicci et al., 2020), the phenotypic features these TRBV4-1 or TRBV7-9-biased cells is less clear. In the case of CD1c-endo and CD1c-PM-reactive cells, however, analysis of memory markers suggest that they may follow adaptive-like dynamics, spanning naïve through different memory subsets(Gherardin et al., 2021a).

Here, we report the detailed cryo-EM structure of CD1c, two lipid ligands and a TCR, along with tetramer staining patterns, activation-based assays with diverse antigens and mutational analysis of functional hotspots on CD1c for recognition of model mycobacterial antigens. These studies begin to answer both general topological questions regarding CD1c antigen display and connect them to specific residues in the public *TRBV7-9* TCR, and because key interactions are generated by conserved residues, suggests a general model of antigen recognition by the larger conserved T cell repertoire. The recently solved structure of the 3C8 autoreactive TCR showed that it bound in a lipid-independent manner. Here structural and functional data show how representative TCRs encoded with the newly described *TRBV7-9* motif have dependence on, and preference for, the PM antigen in favour of MPM, as well as both structural and functional evidence for specific TCR contact with protruding the PM antigenic head group.

Overall, the MPM7 footprint and hotspot analysis are connected to phosphoantigen-dependent responses. Further, reorganisation of the particularly flexible CD1c backbone structure positions the cationic cup of the TCR next to a dominant anionic phosphate of CD1c bound PM. This study shows further that there would be steric clashes between the mannosyl headgroup of MPM and the MPM7 TCR, explaining the preference for PM over MPM by all CD1c TCRs studied here, except CD8-1. Thus, this work provides detailed functional insight combined with direct structural analysis into CD1c-antigen-TRBV7-9^+^ TCR contact.

In summary, we show that the *ex-vivo* use of CD1c tetramers can track antigen-specific CD1c-restricted T cells, paving the way for the study of these cells in disease and clinical settings. We furthermore provide a genetic and molecular basis for the fine specificity and recognition of a clinically-relevant mycobacterial lipid presented by CD1c.

## Materials and Methods

### Human PBMC samples

Most PBMC samples were isolated from healthy donor-derived buffy packs provided by the Australian Red Cross under agreement numbers: 17-08-VIC-16, 18-08-VIC-12, 20-10VIC-14, 22-11VIC-04 and 24-10VIC-15 in accordance with PBMC were isolated by density gradient using Ficoll-Paque Plus (GE Healthcare) and either used fresh or cryopreserved in liquid nitrogen for subsequent use.

Seattle PBMC samples were isolated from healthy human adult donors (Bloodworks, Seattle, WA) by density centrifugation with Ficoll-Paque PLUS media and were cryopreserved for subsequent use. Mycobacteria-exposed PBMC samples were obtained from a subset of 6363 South African adolescents who were enrolled into a study that aimed to determine the incidence and prevalence of tuberculosis infection and disease in South Africa(Mahomed et al., 2011). Twelve-to 18-year-old adolescents were enrolled at eleven high schools in the Worcester region of the Western Cape of South Africa. Subjects were screened for the presence of latent tuberculosis by a TST and IFN-γ release assay (IGRA) QuantiFERON-TB GOLD In-Tube (QFTG) (Cellestis Inc.) at study entry. PBMC were isolated from freshly collected heparinized blood via density centrifugation and cryopreserved. For this work, a sample of six Mtb-infected (TST+) and six Mtb-uninfected (TST-) adolescents were selected based on the availability of PBMC.

The study protocols were approved by either the University of Melbourne Human Ethics Committee (protocols 13000 and 30058) or the IRBs of the University of Washington and the University of Cape Town. Written informed consent was obtained from all adult participants for the South African study samples as well as from the parents and/or legal guardians of the adolescents who participated. In addition, written informed assent was obtained from the adolescents in the South African samples.

### Lipids

Lipids were dissolved by sonication in tris-buffered saline with 0.05% tyloxapol (Sigma-Aldrich) and stored at-20°C prior to use. When required, lipids were thawed and re-sonicated for approximately 30 min or until clear. Loading was performed overnight at room temperature (RT). CD1d was loaded at a 3:1 ratio of lipid:CD1 using KRN7000. Other lipids including synthetic phosphomycoketide (PM; C32) PM, mannosyl phosphomycoketide (MPM; C32)(Buter et al., 2013; van Summeren et al., 2006), mannosyl phosphomycoketide-3 (MPM-3, C32, a difluoromethylene-modified version of MPM that is resistant to hydrolysis(Reijneveld et al., 2021)), ganglioside GD3, sulfatide (C24:1), phosphatidylcholine (PC), phosphatidylinositol (PI), phosphatidic acid (PA), phosphatidylglycerol (PG), phosphatidylethanolamine(PE) and phosphatidylserine (PS) (Avanti Polar Lipids) were loaded at 12:1.

### Flow Cytometry

For PBMC staining, cells were first incubated in 50 nM dasatinib (STEMCELL Technologies 73082) in PBS for 30 min at 37°C. After washing in FACS buffer (PBS supplemented with 2% FBS and 2μg/ml DNAse, Roche 10104159001), cells were then blocked in FACS buffer containing anti-CD36 (10 μg/ml, clone 5-721, Biolegend 336202) and Fc-block (1:50, Miltenyi 130-059-901) for 15 min at 4°C. CD1c and CD1d tetramers (produced in-house as previously described(Gherardin et al., 2021a) made from streptavidin phycoerythrin (PE; Becton Dickinson 554061) were the added directly to the cells at a final concentration of 1 μg/ml and incubated for a further 30 min at RT. Cells were then washed with FACS buffer and stained for 25 min at 4°C in FACS buffer and antibodies and dyes including LIVE/DEAD Fixable Near-IR Dead Cell Dye (1:500, Thermo Fisher Scientific), anti-CD14 APC-Cy7 (1:100, clone MϕP-9, Becton Dickinson 557831), anti-CD19 APC-Cy7 (1:100, clone SJ25C1, Becton Dickinson 557791), anti-CD3 BUV395 (1:50, clone UCHT1, Becton Dickinson 563546), anti-CD4 BUV496 (1:100, clone SK3, Becton Dickinson 612936), anti-CD8 BUV805 (1:300, clone SK1, Becton Dickinson 612889), and anti-TCRγδ PE-Cy7 (1:100, clone 11F2, Becton Dickinson 752023). Cells were then washed once more in FACS buffer, fixed in PBS supplemented with 2% paraformaldehyde (PFA; ProSciTech C004-100) for 10 min at RT. Cells were washed a final time in FACS buffer before being passed through 70 μm mesh and acquired on an LSRFortessa flow cytometer (Becton Dickinson). Cells were gating by first removing cells in the APC-Cy7 channel (dead cells, B cells and monocytes), followed by doublet exclusion using FSC-A versus FSC-H and gating on lymphocytes using FSC-A versus SSC-A. T cells were then defined as CD3^+^ cells. For magnetically enriched cells, after enrichment, cells were then stained as above with additional Fc-block in the antibody cocktail. For FACS sorting post-enrichment as well as analysis of *in vitro*-expanded cells and single cell sorting of *in vitro*-expanded cells, cells were harvested, blocked in FACS buffer with anti-CD36 (10 μg/ml) and Fc-block (1:50) as above, and antibody cocktail as above added directly to the block with the addition of 1 μg/ml CD1 tetramer. Cells were incubated for 30 min at 4°C, washed twice in FACS buffer and then passed through 70 μM mesh before FACS-sorting on an ARIA III flow cytometer (Becton Dickinson) with gating as per PBMC analysis above. For primary T cell activation assays, co-cultures were harvested, washed in FACS buffer and then stained in FACS buffer with LIVE/DEAD Fixable Near-IR Dead Cell Dye (1:500), anti-CD69 PE-Cy7 (1:100, clone FN50, Becton Dickinson 557745) and anti-CD25 APC (1:100, clone BC96, Biolegend 961721). Cells were then washed in FACS buffer, fixed in 2% PFA as above, washed once more in FACS buffer and then filtered through 70 μm mesh prior to acquisition on an LSRFortessa (Becton Dickinson). For analysis, dead cells were first excluded removing viability-dye positive cells, followed by double exclusion and FSC-A versus SSC-A gating as above. CTV-labelled K562 cells were then removed, and T cells assessed for CD69 or CD25 expression. For transient transfections, HEK293T.SCARB1^-/-^ cells(Gherardin et al., 2021a) were mechanically harvested, and passed through 100 μm mesh. After pelleting, cells were resuspended and incubated for 30 min at RT in FACS buffer supplemented with antibodies and dyes including LIVE/DEAD Fixable Near-IR Dead Cell Dye (1:500, Thermo Fisher Scientific), anti-CD3 BUV395 (1:100, clone UCHT1, Becton Dickinson 563546) and CD1 tetramers at 1 μg/ml, generated using streptavidin-PE (Becton Dickinson 554061). Cells were then washed in FACS buffer, fixed as above and washed once more with FACS buffer prior to acquisition on an LSRFortessa flow cytometer (Becton Dickinson). Cells were first gated by removing viability dye positive cells, following by doublet exclusion and FSC-A and SSC-A gating as above. Cells were then gated on GFP+ cells and assessed for CD3 versus tetramer. A gate was set on a consistent window of CD3 between stains for a given TCR, and the median-fluorescence intensity (MFI) of tetramer-PE staining determined for cells falling within that gate. For SKW-3 co-culture experiments, co-cultures were harvested, washed in FACS buffer and then stained for 30 min at RT with FACS buffer supplemented with LIVE/DEAD Fixable Near-IR Dead Cell Dye (1:500, Thermo Fisher Scientific), anti-CD3 BUV395 (1:100, clone UCHT1, Becton Dickinson 563546) and anti-CD69 PE-Cy7 (1:100, clone FN50, Becton Dickinson 557745). Cells were then washed in FACS buffer, fixed in 2% PFA as above and washed once more in FACS buffer prior to passing through 70% mesh and acquisition on a LSRFortessa Flow Cytometer (Becton Coulter). Cells were first gated on viability dye negative cells followed by doublet exclusion as above and FSC-A and SSC-A gating. Cell Trace Violet positive cells (APCs) were then removed and GFP^high^ cells gated prior to assessing the MFI of CD69 PE-Cy7. All flow cytometry data was analysed using FlowJo Software version 10.10 (Becton Dickinson).

For PBMCs from Seattle and South Africa, cells were stained with LD Aqua (1:200, Thermo Fisher Scientific, L34966) for 15 minutes at room temperature, then washed with PBS. Cells were then incubated in 50 nM dasatinib (STEMCELL Technologies 73082) in PBS for 30 min at 37°C. Cells were then directly blocked in PBS containing anti-CD36 (10 μg/ml), Fc-block, and 10% human serum (Valley Biomedical HS1004CHI) for 15 min at RT. CD1c tetramers (loaded in-house as previously described(Layton et al., 2021), using biotinylated CD1c monomers from the National Institutes of Health (NIH) Tetramer Core Facility; Emory University, Atlanta, GA) were generated from streptavidin APC (Thermo Fisher Scientific, S868) were added directly to the cells at 1:50 and incubated for a further 1 hour at RT. Cells were then washed with FACS buffer and stained for 30 min at 4°C in FACS buffer and antibodies including anti-CD3 ECD (1:100, clone UCHT1, Beckman Coulter IM2705U), anti-TCRγδ PE Vio770 (1:50, clone 11F2, Miltenyi 130-113-505), and anti-Vδ2 AF700 (1:400, clone B6, BioLegend 331416). Cells were then washed once more in FACS buffer, again in PBS, then fixed in PBS supplemented with 1% paraformaldehyde (PFA; Electron Microscopy Sciences, 15712-S) for 15 min at 4°C. Cells were washed a final time in PBS buffer before being resuspended in 2mM EDTA (Thermo Fisher Scientific 15575020) in PBS and acquired on an LSRFortessa flow cytometer (Becton Dickinson). Cells were first gated on lymphocytes using FSC-A versus SSC-A, then doublet exclusion using FSC-A versus FSC-H, then gating out dead LD Aqua^+^ cells, and T cells were then defined as CD3^+^ cells. Tetramer gates were set using no tetramer controls as reference.

### Magnetic enrichments

PBMCs were isolated from buffy packs as above. 150 million cells per sample were incubated with 50 nM dasatinib (STEMCELL Technologies 73082) in PBS for 30 min at 37°C. Cells were then blocked in FACS buffer containing anti-CD36 (5 μg/ml, clone 5-721, Biolegend 336202) for 15 min at 4°C. CD1c tetramers labelled made from streptavidin phycoerythrin (PE; Becton Dickinson 554061) were the added directly to the cells at a final concentration of 1 μg/ml and incubated for a further 30 min at room temperature (RT). Cells were then washed three times in FACS buffer before being resuspended in 1200 μl MACS buffer (PBS supplemented with 0.5% FBS with 5 mM EDTA) with 150 μl anti-PE MicroBeads (Miltenyi 130-048-801) and incubated for 20 min at 4°C. Cells were then washed twice in MACS buffer before being resuspended in 3 mL MACS buffer and passed through a 70 μM SmartStrainer (Miltenyi 130-110-916) and run over a magnetic LS Column (Miltenyi 130-042-401). Flow through was collected and passed over the column once more. Columns were then washed with 12 mL of MACS buffer before being removed from the column and flushed with 5 mL MACS buffer.

### In vitro T cell expansion

Magnetically-enriched cells were cultured overnight in complete media (RPMI-1640 supplemented with 10% fetal bovine serum (FBS) (JRH Biosciences), penicillin (100 U/ml), streptomycin (100 μg/ml), GlutaMAX (2 mM), sodium pyruvate (1 mM), nonessential amino acids (0.1 mM), Hepes buffer (15 mM) (pH 7.2 to 7.5) (Invitrogen, Life Technologies), and 2 mercaptoethanol (50 μM, Sigma-Aldrich)) supplemented with 50 U/mL recombinant human IL-2 (Peprotech). Cells were then harvested, stained as above and FACS-Sorted using an ARIA III (Becton Dickinson). Cells were sorted directly into sorted in bulk directly into U-bottom 96-well plates containing T cell media (complete media supplemented with 200 U/mL recombinant human IL-2 (Peprotech), 25 μg/mL recombinant human IL-7 (eBiosciences) and 50 μg/mL recombinant human IL-15 (eBiosciences)) with plate-bound anti-CD3 (10 μg/mL, clone OKT3, Becton Dickinson) plate-bound anti-CD28 (2 μg/mL, clone CD28.2, Becton Dickinson) and soluble phytohemagglutinin (1 μg/mL, Thermo Fisher Scientific). Cells were incubated for 72 hrs at 37°C in 5% CO_2_ before being removed from anti-CD3, anti-CD28 and PHA and placed in fresh T cell media. Cells were cultured for up to 3 weeks, with media being replaced or cells split across multiple wells as the media colour changed or the cells overgrew the wells. Cells were either analysed fresh or cryopreserved in liquid nitrogen for future use.

### Primary T cell activation assays

Wildtype or CD1c overexpressing K562 cells were labelled with Cell Trace Violet (Thermo Fisher Scientific, C34557) as per manufacturer’s instructions. Cells were then incubated for 20 hrs at 37°C, 5% CO_2_ in 96-well U-bottom plates (Corning) with *in vitro* expanded polyclonal T cells with or without exogenous PM lipid. Some wells contained T cells alone, or T cells plus Human T-Activator CD3/CD28 Dynabeads (Thermo Fisher Scientific). Cells were then harvested and stained as above.

### Single cell T cell receptor sequencing

T cell receptor sequencing was performed as previously described(Dash et al., 2011). In brief, single αβ T cells were sorted in 96-well PCR plates (Thermo Fisher Scientific), capped, and frozen at-80°C for future use. For sequencing, cDNA was first generated using superscript VILO cDNA synthesis kit (Thermo Fisher Scientific 11754250) before cDNA was used in two rounds of semi-nested PCR with primers sets for αβTCR genes using 2X PCR Mastermix (Promega M7505). PCR products were analysed by Gel electrophoresis and where PCR product of appropriate size was evident, the product was treated with ExoSAP-IT PCR Product Cleanup Reagent (Thermo Fisher Scientific (78201.1.ML) before Sanger Sequencing using the internal reverse primers. Sequences were analysed using the IMGT V-QUEST tool.

### Generation of TCR and CD1-encoding plasmids

dsDNA corresponding to full-length TCR-α genes linked to full-length TCR-β genes via a T2A-linker or the full-length CD1c gene was flanked with EcoRI and XhoI restriction enzyme sites and synthesised (Integrated DNA Technologies). DNA was then cloned into pMIG II plasmid (Addgene 52107) for subsequent use in transient transfection or retroviral transduction experiments.

### Transient transfections

Transient transfections were performed as previously described(Gherardin et al., 2016). In brief, pMIG II plasmids encoding full-length αβ TCR were co-transfected using FUGENE HD transfection reagent (Promega, E2311) into SCARB1^-/-^ 293T cells(Gherardin et al., 2021a) with a pMIG II plasmid encoding all CD3ε, δ, γ and ξ subunits linked by P2A linkers. Transfected cells were incubated for 48 hrs prior to harvesting and staining as per above.

### Generation of K562 and SKW-3 cell lines

To generate CD1c overexpressing K562 cells and TCR-expressing SKW-3 cells, retroviral transduction was used as previously described(Holst et al., 2006). pMIG II plasmids described above were used to generate retrovirus with HEK293T cells used as packaging cell lines. K562 or SKW-3 cells were incubated in fresh retrovirus twice daily for 4 days, prior to further culture in complete media for one week and subsequent FACS-sorting for GFP^high^ cells. Cells were then expanded and cryopreserved in liquid nitrogen for further use.

### SKW-3 activation assays

Antigen-presenting cells (K562 cells or C1R cells including CD1c-overexpressing C1R cells and mutants thereof, as previously described(Wun et al., 2018) were first labelled with Cell Trace Violet (Thermo Fisher Scientific, C34557) as per manufacturer’s instructions. Cells were then co-incubated in the presence or absence of SKW-3 T cells with or without exogenous PM or MPM. Cells were cultured for approximately 20 hrs before harvesting and staining for flow cytometric analysis as described above.

### Protein expression, purification, and lipid loading

Genes encoding human CD1c and human β2-microglobulin (β2m) were cloned into a single construct separated by a 2A self-cleavage peptide in the pHLSEC vector, with the fos/jun leucine zippers, Igκ leading sequence, a 6×His-tag and an Avi-tag, using the same design as in our previous study(Cao et al., 2025). The single-plasmid construct was transfected into Expi293F GntI(-) cells. The secreted CD1c-β2m heterodimer complex, referred hereafter as CD1c, was pulled down by nickel affinity chromatography, then subjected to size-exclusion chromatography (HiLoad Superdex 200 16/60, Cytiva). The same construct design and purification procedure were used for human CD1d (hCD1d), which was used as reference sample for surface plasmon resonance experiment. Purification was validated by SDS-PAGE. PM was loaded onto CD1c by incubating the protein at a molar ratio of 12:1 at room temperature in the presence of 0.05% tyloxapol. GD3 was further loaded onto CD1c-PM by incubating at a molar ratio of 12:1 at room temperature in the presence of 0.05% tyloxapol. The residual lipids and tyloxapol were then removed using MonoQ anion exchange chromatography (Cytiva). Loading efficiency was validated using Novex iso-electric focusing (Thermo Fisher Scientific).

Genes encoding α-and β-chains of MPM7 TCR (wildtype or mutants) were cloned separately into pET30a vector. The separate plasmids were transformed into *E. coli* BL21 (DE3) cells. Protein expression was induced using 0.5 mM isopropyl β-D-1-thiogalactopyranoside (I2481C25, GoldBio). After 4 hours at 37 °C, cells were harvested then lysed with 1 mg/mL lysozyme supplemented with 0.5 mg/mL DNase I and 2 mM magnesium chloride. Inclusion bodies were washed a buffer containing 50 mM Tris, 100 mM NaCl, 1 % (v/v) Triton X-100, 2 mM dithiothreitol (DTT), 1 mM EDTA, and 0.5 mM phenylmethylsulfonyl fluoride (PMSF, P-470-50, GoldBio), then solubilized in a buffer containing 50 mM Tris, 6 M guanidine hydrochloride, and 10 mM DTT at pH 8.0. Fifty mg of each chain were gradually injected into a refolding buffer comprising 50 mM Tris, 5 M urea, 440 mM L-arginine, 4 mM EDTA, 0.5 mM PMSF, 2 mM reduced glutathione, and 0.2 mM oxidised glutathione at pH 8.0. After incubation at 4°C for 48 h, the mixture was dialysed against a buffer containing 10 mM Tris pH 8.0 and subjected to sequential DEAE-cellulose anion exchange, strong anion exchange using HiTrapQ, size-exclusion chromatography using HiLoad 16/600 Superdex 200 pg column, and hydrophobic interaction chromatography using HiTrap Phenyl column (Cytiva). Purification was validated by SDS-PAGE.

### Surface plasmon resonance

CD1c and hCD1d that contained endogenous lipids acquired from the cultured mammalian cells (CD1c-endo and hCD1d-endo) were biotin labelled using the BirA ligase enzyme. PM and GD3 were further loaded onto CD1c as described in the preceding section. A total of 2500 units of each CD1-lipid complex was coupled onto a streptavidin sensor chip SA (Cytiva), with hCD1d-endo immobilised at the first flow-cell as the reference control. Steady-state equilibrium affinity of MPM7 TCR and its indicated mutants was measured by passing the TCRs over the sensor chip at 6 μL/min for 120 s, using a buffer comprising 10 mM HEPES and 150 mM NaCl, pH 7.4. Serial dilutions of the TCRs were injected at the concentrations of 10, 5, 2.5, 1.25, 0.625, 0.3125, 0.15625 and 0 μM. Affinity values and sensorgram plots were analysed and generated using GraphPad Prism software.

### Grid preparation, cryo-EM data acquisition, image processing and model building

CD1c loaded with PM and GD3 (CD1c-PM-GD3) was incubated with the MPM7 TCR at a molar ratio of 1.1:1 in a buffer comprising 10 mM HEPES and 150 mM NaCl, pH 7.4, for 16 hours at 4 °C. The complexed fractions were pooled by size-exclusion chromatography (Superdex 200 Increased 10/300 GL column, Cytiva), at a flow rate of 0.2 mL/min. The sample at 2 mg/mL was blotted onto Quantifoil R1.2/1.3 grids on 200-mesh copper (Quantifoil Micro Tools) employing a Vitrobot Mark IV (Thermofisher Scientific) with zero blotting force for 6 s at 4 °C and 100% humidity, with drainage for 1 s prior to plunging into liquid ethane. The grids were transferred to a Titan Krios transmission electron microscope (Thermofisher Scientific) equipped with a Gatan K3 direct detection camera and a Gatan Quantum energy filter. Data acquisition was carried out at 300 kV, with a 50-μm C2 aperture at a nominal magnification of 105,000x, a pixel size of 0.82 Å, a spherical aberration of 2.7 mm, a total exposure dose of 60 eÅ^-2^, and a defocus range of-0.5 to-1.5 μm. A total of 10,422 micrographs were collected.

Images were calibrated through gain-reference correction using Relion(Scheres, 2012). Further processing was undertaken in CryoSPARC(Punjani et al., 2017). After patch motion correction and patch CTF estimation at a maximal resolution of 3 Å, the movie set was split into 521-exposure groups, followed by blob picking with minimum and maximum particle diameters of 80 and 200 Å, respectively, on a randomly combined dataset of 1,563 exposures. An initial set of 977,212 particles, extracted with a box size of 280 px, underwent multiple rounds of 2D classification, narrowing the dataset to 33,151 particles, which subsequently served as a template for picking 6,105,016 particles from the overall dataset. The newly curated particle set again underwent multiple rounds of 2D classification, yielding 105,927 particles, from which the *ab initio* reconstruction further refined the dataset to a reasonable volume from 20,082 particles (18.9%). Subsequent non-uniform (NU) refinement resolved the volume to 4.04 Å. This particle set was then subjected to orientation rebalancing to reduce orientation bias, and the resulting particle set served as template for deep picking using Topaz(Bepler et al., 2019) on the overall dataset. A set of 566,251 particles, extracted with a box size of 300 px, was subjected to one round of 2D classification, narrowing the dataset to 514,401 particles. A three-class *ab initio* reconstruction identified a reasonable volume from 205,371 particles (39.9% of the curated 514,401 particle set), which was subsequently resolved to 3.28 Å via NU-refinement. This volume served as the reference for one round of heterogenous refinement, improving the resolution of the volume to 3.12 Å from 131,430 particles (37.7%). This newly cleaned particle set was subjected to a sequential series of reference-based motion correction, global CTF refinement, NU-refinement, and local refinement, yielding a final resolution of 2.90 Å from 130,855 particles (25.4%). The 42-day workflow is summarized in **Figure S5**.

A search model of the MPM7 TCR was generated in Swiss-Model(Waterhouse et al., 2018), while the PDB entry 6C15(Wun et al., 2018) was used as the search model for CD1c. Models were placed into the 2.90 Å resolution density map using the phenix.dock_in_map tool(Liebschner et al., 2019) prior to manual correction in Coot(Emsley et al., 2010). The final model was refined using Servalcat via CCP-EM suite(Wood et al., 2015; Yamashita et al., 2021) and phenix.real_space_refine tool(Liebschner et al., 2019).

### Data presentation and statistical analysis

Graphs were generated using PRISM software (version 10, Graphpad) and statistical analysis performed within PRISM. Where statistical analysis was performed, specific statistical tests are outlined in the figure legends. Flow cytometry plots were generated using Flowjo version 10.10 (Becton Dickinson). Structural illustrations were rendered in PyMol and ChimeraX. All figures were generated Adobe Illustrator 2025.

## Acknowledgements

We thank staff from the flow cytometry facilities at the Department of Microbiology and Immunology at the Peter Doherty Institute. We thank the Monash Ramaciotti Centre for Cryo-Electron Microscopy, a Node of Microscopy Australia, for the use of instruments and assistance. This work was supported by the Australian Research Council (DP210103064) to DIG and NAG. NAG/AS were supported by ARC Discovery Early Career Awards (DE21010070 and DE210101031) and now NHMRC Emerging Leadership Grants (2027058 and 2027104). DIG and JR were supported by NHMRC Investigator Awards (2008913 and 2008981). CS was supported by the US National Institutes of Health (R01-AI146072)

## Author contributions

NAG, DIG, AS and JR conceptualised, planned and managed the project. T-PC, CaS, SJR, TT, JK, HV, QG, APU, AS, TJS, ChS and NAG performed the experiments and analysed the data. DBM provided intellectual input and critical reagents. CS undertook the cellular immunology-based experiments. T-PC designed and performed surface plasmon resonance experiments, cryo-EM sample preparation, image processing and structure solving, and analysed the data. JK and QG assisted with research. HV performed advanced cryo-EM data collection. NAG, T-PC, AS, JR and DIG drafted the manuscript. AS and JR conceptualised the structural and biophysical work, supervision, project management/ funding. All authors reviewed and contributed to the manuscript.

## Supplementary information

**Figure S1:**
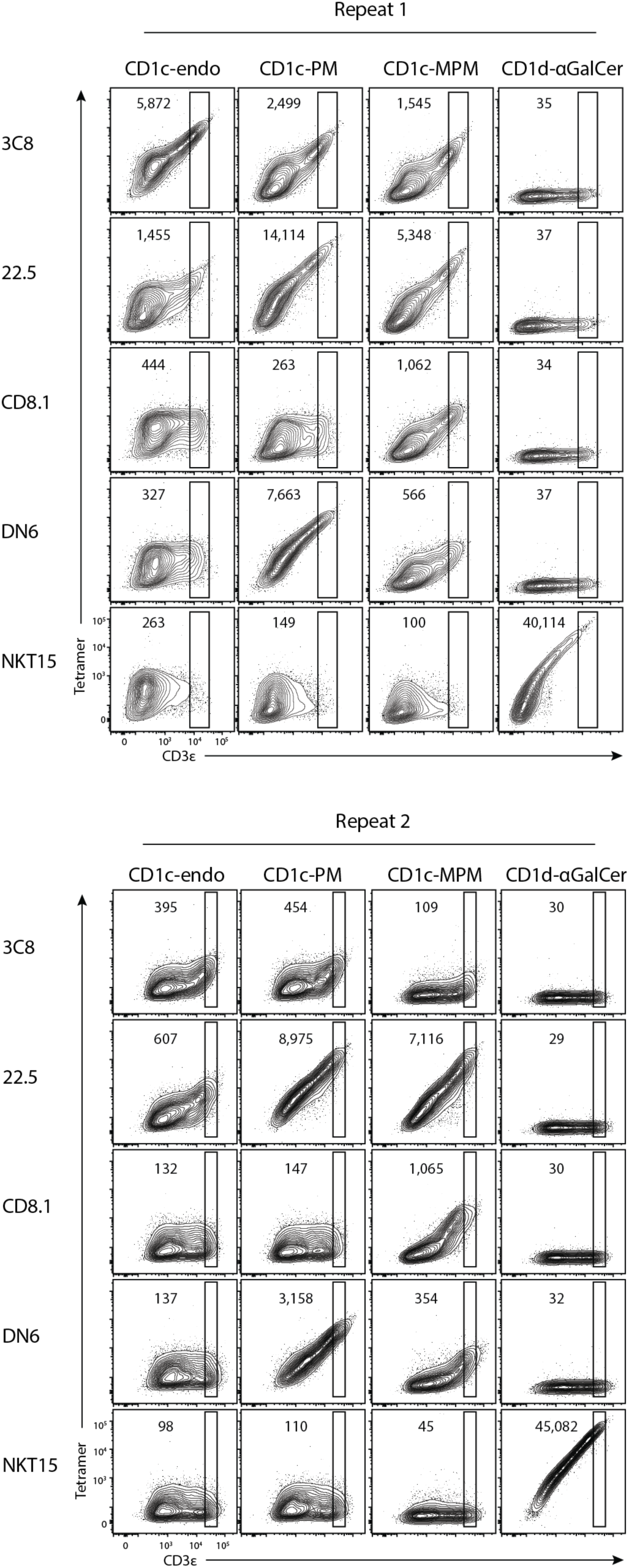
Repeat CD1c tetramer staining for. Figure 1C. Flow cytometric contour plots showing CD1 tetramer staining on HEK293T.SCARB1^-/-^ cells transiently transfected to express CD1-restricted TCRs. Numbers show MFI of gated population which is consistently placed within a TCR-transfected cell line. Data is a repeat of the experimental data shown in Figure 1B.

**Figure S2:**
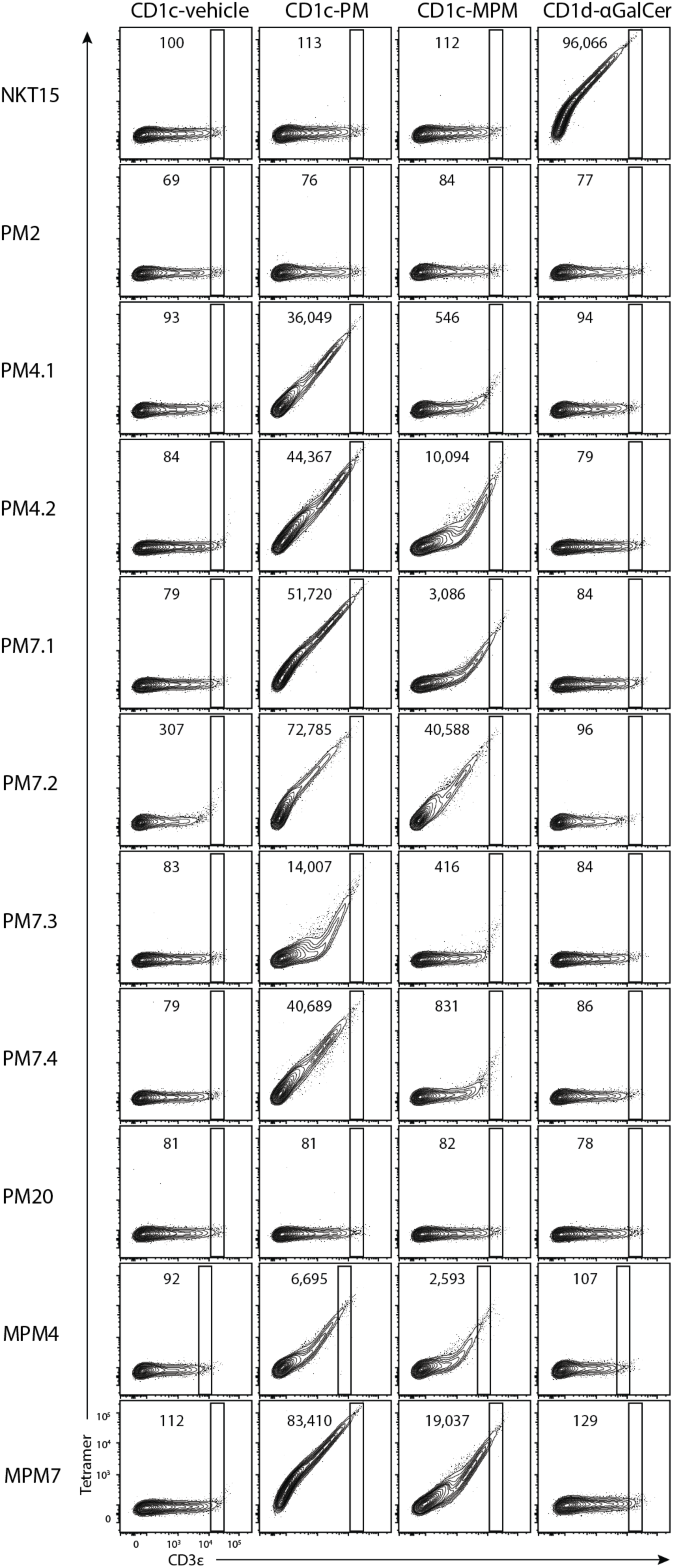
Repeat CD1c tetramer staining for. Figure 3B. Flow cytometric contour plots showing CD1 tetramer staining on HEK293T.SCARB1^-/-^ cells transiently transfected to express CD1-restricted TCRs. Numbers show MFI of gated population which is consistently placed within a TCR-transfected cell line. Data is a repeat of the experimental data shown in Figure 3B.

**Figure S3:**
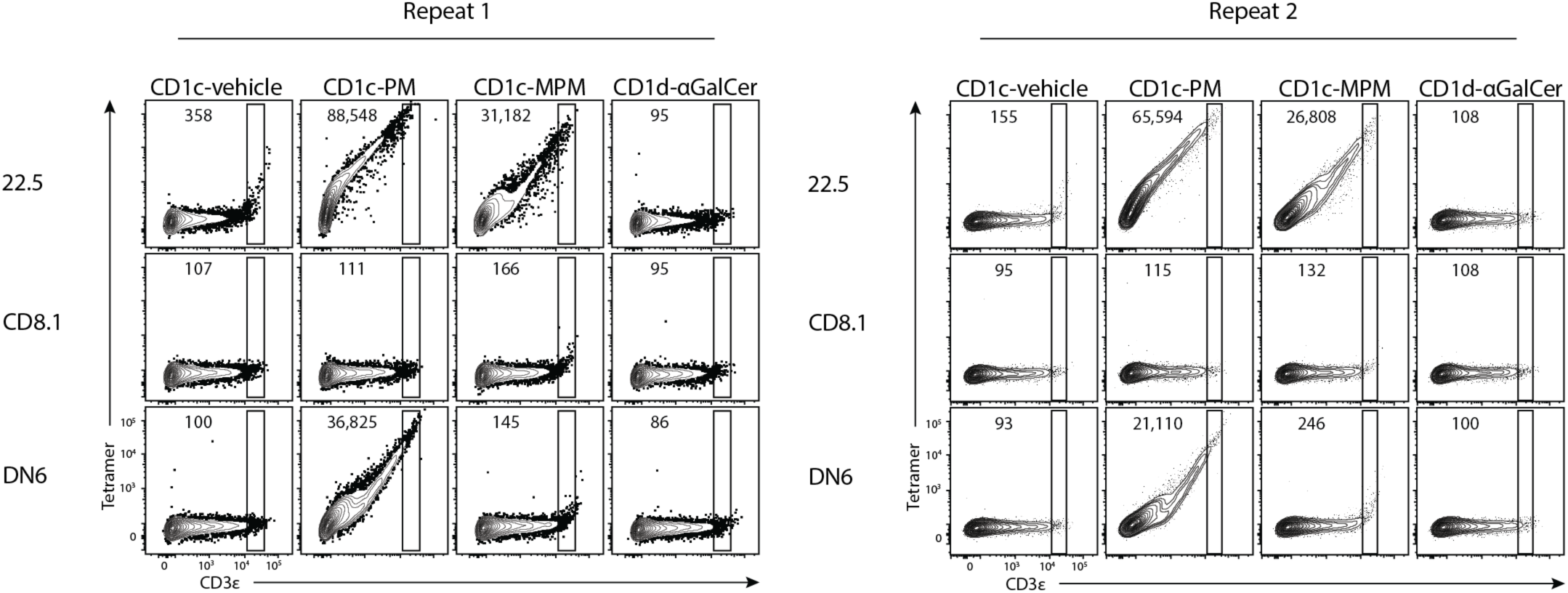
Control TCR CD1c tetramer staining. Flow cytometric contour plots showing CD1 tetramer staining on HEK293T.SCARB1^-/-^ cells transiently transfected to express CD1-restricted TCRs and stained in parallel to those stained in Figures 3B and 4B. Numbers show MFI of gated population which is consistently placed within a TCR-transfected cell line. Data is depicted from n=2 independent experiments.

**Figure S4:**
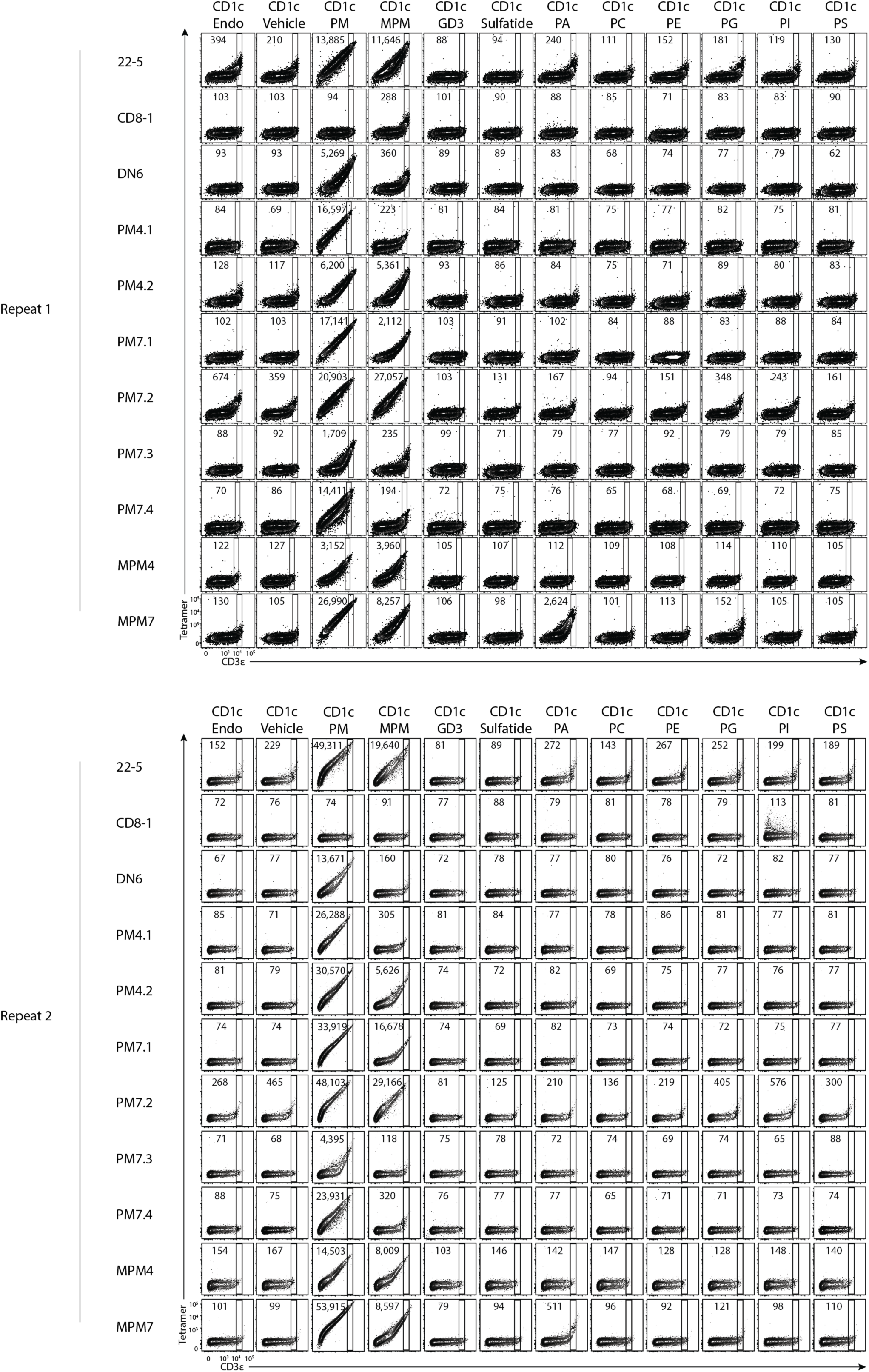
Antigen-specificity of CD1c-PM-restricted TCRs. Flow cytometric contour plots showing CD1 tetramer staining on HEK293T.SCARB1^-/-^ cells transiently transfected to express CD1-restricted TCRs. Numbers show MFI of gated population which is consistently placed within a TCR-transfected cell line. Data is depicted from n=2 independent experiments.

**Figure S5.**
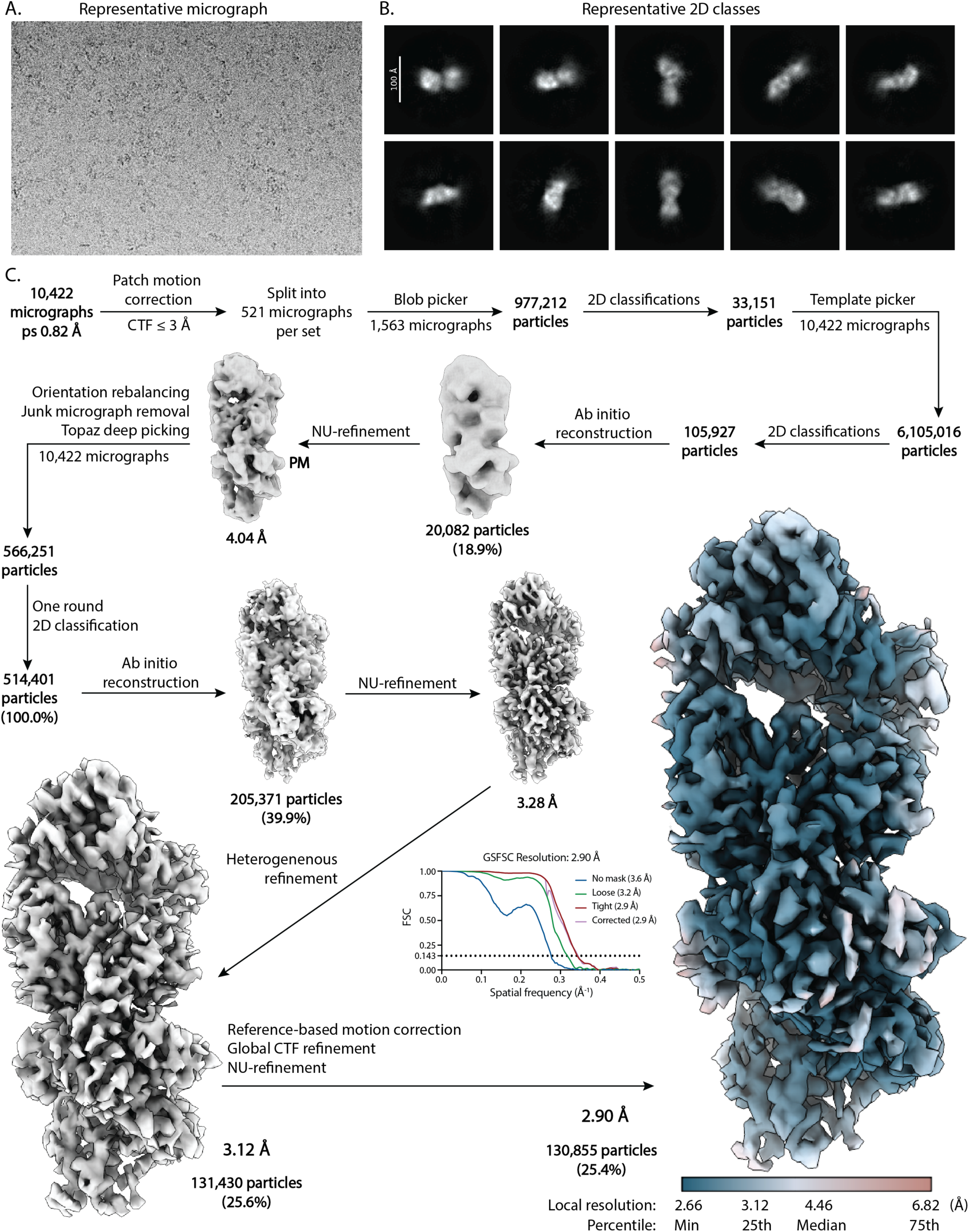
42-day Cryo-EM workflow. **A.** Representative micrograph from a total of 10,422 micrographs. **B.** Representative ten 2D classes from a total of 45 classes of 514,401 particles deep-picked by Topaz. **C.** Processing tree for determining the 2.90 Å resolution structure of MPM7 TCR-CD1c-PM-GD3.

**Figure S6.**
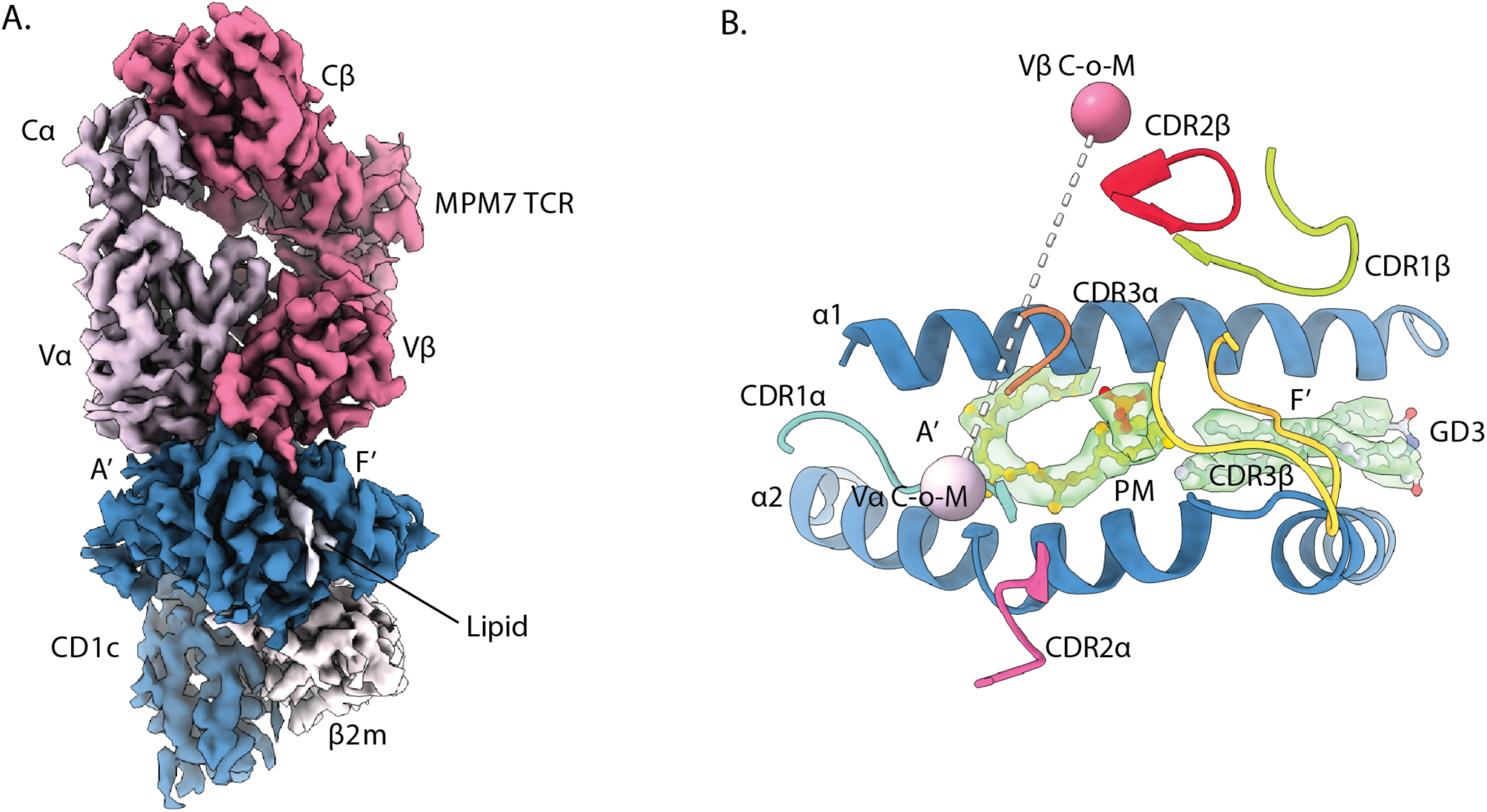
Overall cryo-EM structure of MPM7 TCR-CD1c-PM-GD3 complex. **A.** Cryo-EM density of the MPM7 TCR-CD1c-PM-GD3 complex. The α-chain of MPM7 TCR is coloured light-violet and the β-chain is coloured pink. CD1c and β2m is coloured steel-blue and white, respectively.**B.** Electron density of PM and GD3 in the A’-and F’-pockets of CD1c, respectively. Positions of the CDR1, CDR2, and CDR3 loops of the MPM7 TCR are indicated. Centre-of-mass (C-o-M) of Vα and Vβ are marked as spheres.

**Figure S7.**
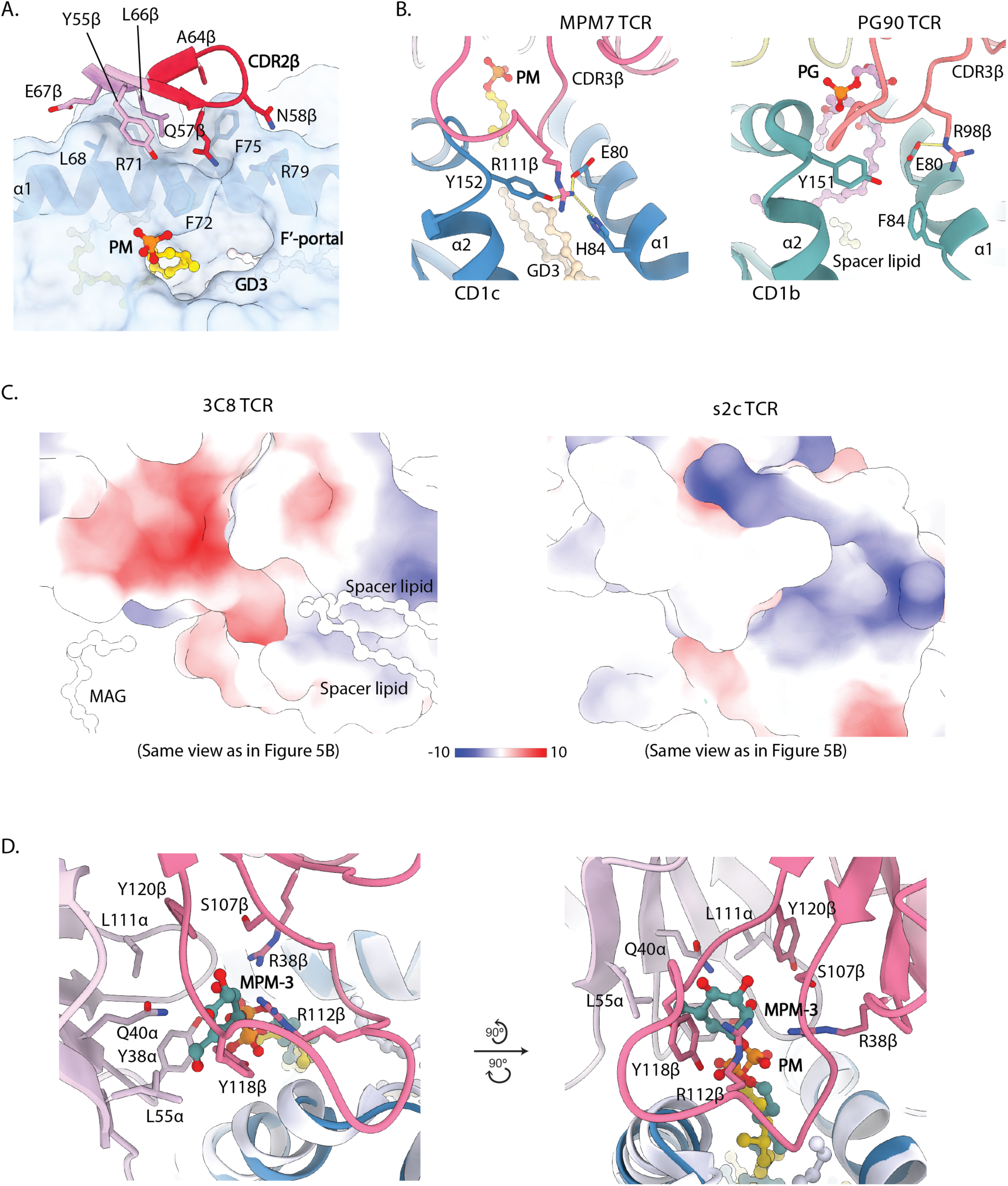
Structural features of MPM7 TCR-CD1c-PM comparing to other structures. **A.** Details of the interaction between CDR2β and adjacent FWβ regions of MPM7 TCR to CD1c. CD1c is displayed as surface representation. **B.** Left, Details of interaction between R111^β^ of MPM7 TCR and the E80-Y152-H84 cluster of CD1c. Right, Details of interaction between R98^β^ of PG90 TCR and the E80-Y151-F84 cluster of CD1b (PDB ID: 5WKI). **C.** Electrostatic surface of 3C8 TCR (left) and s2c TCR (right) that binding to CD1c, in the same view as in Figure 5B. **D.** Details of residues at the TCR-CD1c interface that create a steric hindrance preventing the recognition of the mannose headgroup. The colour scheme for the MPM7 TCR-CD1c-PM complex is the same as in Figure 6. The binary structure of CD1c presenting MPM-3 analogue (PDB ID: 7MX4) is coloured in white, and the MPM-3 analogue is shown in teal.

**Figure S8.**
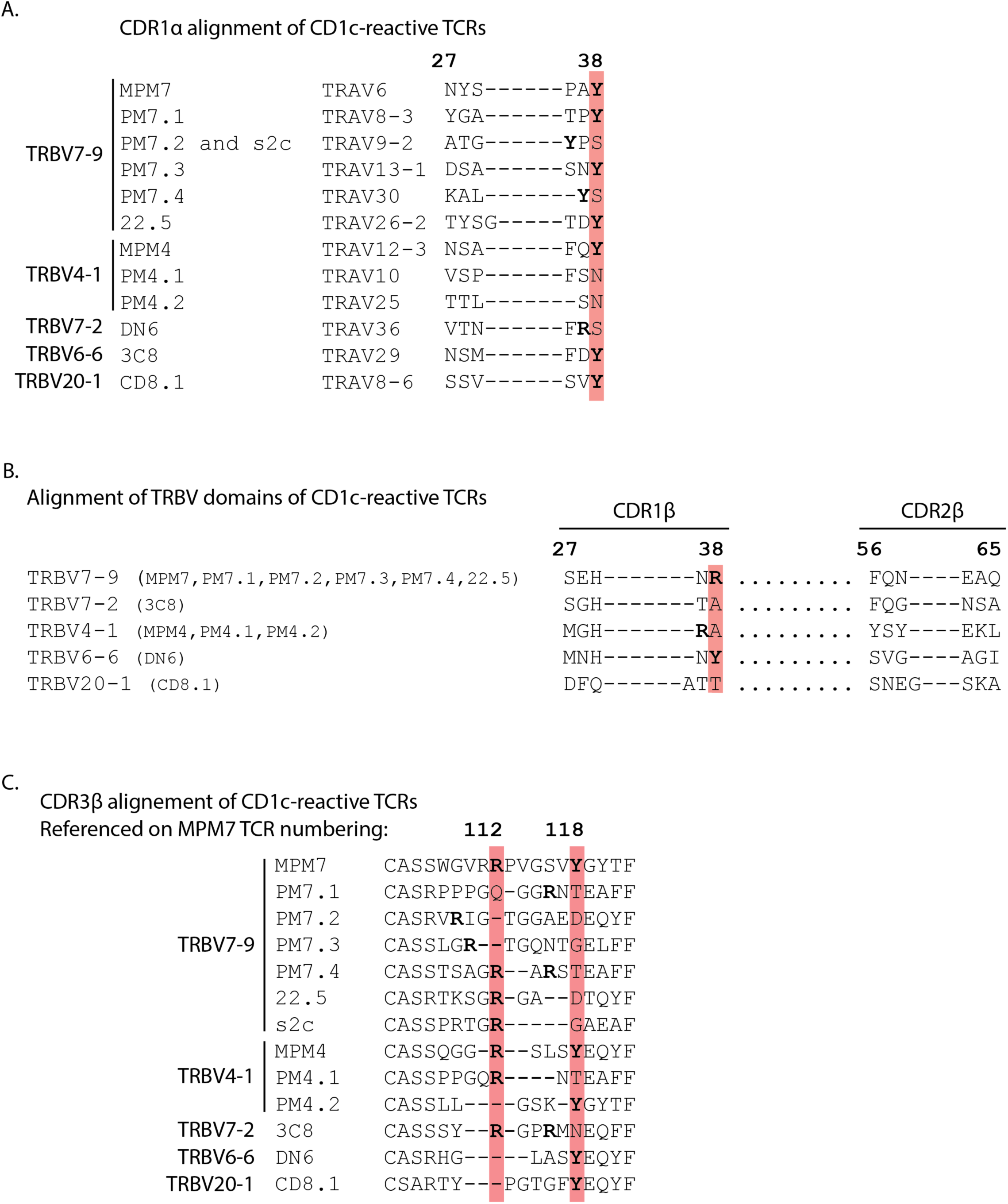
Sequence alignment of CD1c-lipid-reactive TCRs. **A.** Comparison of CDR1α regions of CD1c-restricted TCRs using diverse TCR-α chains **B.** Comparison of CDR2β regions of CD1c-restricted TCRs using diverse TCR-β chains **C.** Comparison of CDR3β regions of CD1c-restricted TCRs using diverse TCR-β chains.

---

**Table S1.** Patient cohort characteristics.

| **Cohort** | **Sample #** | **QFTG** | **TST** | **TSTResult** | **Age** | **Sex** | **Ethnicity** | **Household TB contact** |
| --- | --- | --- | --- | --- | --- | --- | --- | --- |
| *ACS TST+* | 01-0774 | Pos | Pos | 15 | 12 | female | black | n |
| *ACS TST+* | 09-0428 | Pos | Pos | 17 | 13 | female | coloured | n |
| *ACS TST+* | 03-0709 | Pos | Pos | 16 | 14 | female | black | y |
| *ACS TST+* | 09-0043 | Pos | Pos | 17 | 16 | female | coloured | n |
| *ACS TST+* | 01-0272 | Pos | Pos | 12.4 | 14 | male | coloured | n |
| *ACS TST+* | 02-0156 | Pos | Pos | 14 | 16 | female | coloured | n |
| *ACS TST-* | 01-0676 | Neg | Neg | 0 | 15 | male | coloured | n |
| *ACS TST-* | 01-0913 | Neg | Neg | 0 | 18 | male | coloured | n |
| *ACS TST-* | 09-0107 | Neg | Neg | 0 | 12 | male | black | n |
| *ACS TST-* | 02-0184 | Neg | Neg | 0 | 17 | male | coloured | n |
| *ACS TST-* | 11-0178 | Neg | Neg | 0 | 13 | female | coloured | n |
| *ACS TST-* | 02-0320 | Neg | Neg | 0 | 13 | male | coloured | n |

**Table S2.** TCR sequences from CD1c-PM tetramer^+^ αβ T cells.

| *TRAV* | CDR3α | *TRAJ* | *TRBV* | CDR3β | *TRBJ* | INTENSITY | Co-R | DONOR |
| --- | --- | --- | --- | --- | --- | --- | --- | --- |
| 38-2 | CAKTPNGAQKLVF | 54 | 2 | CASRSLAQGKGYTF | 1-2 | moderate | CD4 | 1 |
| 6 | CALPVLYNQGGKLIF | 23 | 4-1 | CASSQGGGRNQPHF | 1-5 | weak | - | 8 |
| 8-3 | CAVGALSSGGSNYKLTF | 53 | 4-1 | CASSQLPRGGRKDSKEQYF | 2-7 | moderate | - | 7 |
| 10 | CVVRLRNDYKLSF | 20 | 4-1 | CASSPPGQRNTEAFF | 1-1 | moderate | DN | 2 |
| 17 | CATVPTNAGGTSYGKLTf | 52 | 4-1 | CASSLSIWTGRREQYF | 2-7 | weak | CD4 | 3 |
| 19 | CALSDFYGNRLAF | 7 | 4-1 | CASSQAGQRSGYGYTF | 1-2 | moderate | CD4 | 3 |
| 25 | CAGPNYPYAGGTSYGKLTf | 52 | 4-1 | CASSLLGSKYGYTF | 1-2 | moderate | CD4 | 3 |
| 26-2 | CILRIQGAQKLVF | 54 | 4-1 | CASKSAGGSRGYTF | 1-2 | strong | - | 7 |
| 6 | CALNTNAGKSTF | 27 | 5-4** | CASRKTGDNYGYTF | 1-2 | moderate | - | 8 |
| 26-1 | CIVSTLSDGQNFVF | 26 | 5-4** | CASRTRKGGAKTQYF | 2-5 | weak | - | 8 |
| 36 | CAVEARSGATNKLIF | 32 | 5-4** | CASSWLGGGRNEQYF | 2-7 | weak | - | 7 |
| 1-2 | CAVMDSNYQLIW | 33 | 6-4 | CASSDGSQAYNEQFF | 2-1 | weak | CD4 | 2 |
| 21 | CAVIVLGSARQLTF | 22 | 7-2 | CASSLNHAVRQGGTDTQYF | 2-3 | weak | - | 7 |
| 13-2 | CAAXTSYGKLTf | 52 | 7-8 | CASSPGRRVHQPHF | 1-5 | weak | CD8 | 1 |
| 17 | CXTRRDSWGKLQF | 24 | 7-8 | CASRPREGHEQFF | 2-1 | weak | - | 8 |
| 25 | CAGKRRITGGGNKLTF | 10 | 7-8 | CASSLARGFEGVGTEAFF | 1-1 | moderate | DN | 6 |
| 26-1 | CIVRVAGFQKLVF | 8 | 7-8 | CASSLARTGNTEAFF | 1-1 | weak | - | 8 |
| 35 | CAVARGATNKLIF | 32 | 7-8 | CASSPLRVGNQPHF | 1-5 | moderate | CD4 | 3 |
| 36 | CAARRQGAQKLVF | 54 | 7-8 | CASSXMRQGGNQPHF | 1-5 | weak | CD4 | 4 |
| 1-2 | CAVRPGYSSASKIIF | 3 | 7-9 | CASSNRRTSGGYNEQFF | 2-1 | moderate | CD4 | 3 |
| 8-3 | CAVVLTGGGNKLTF | 10 | 7-9 | CASRPPPGQGGRNTEAFF | 1-1 | strong | CD4 | 4 |
| 8-6 | CAVKRGSTLGRLYF | 18 | 7-9 | CASSFFTPTSGSRRSYNEQFF | 2-1 | moderate | CD4 | 3 |
| 9-2 | CALRRRALTF | 5 | 7-9 | CASRVGRTGGAEDEQYF | 2-7 | strong | CD4 | 3 |
| 9-2 | CALRGNEKLTF | 48 | 7-9 | CASRVGHYRTSGGGFGNEQFF | 2-1 | weak | CD4 | 2 |
| 9-2 | CALSAQGAQKLVF | 54 | 7-9 | CASSPHFRVPNNEKLFF | 1-4 | weak | - | 8 |
| 13-1 | CAASTYTGRRALTF | 5 | 7-9 | CASSLGRTGQNTGELFF | 2-2 | moderate | - | 8 |
| 14 | CAMRXIIQGAQKLVF | 54 | 7-9 | CASSAVSRKKPTRGNTIYF | 1-3 | moderate | CD4 | 4 |
| 19 | CALRPTGGGNKLTF | 10 | 7-9 | CASSPDRIWTKNIQYF | 2-4 | weak | CD4 | 9 |
| 20 | CAVRTISGGYNKLIF | 4 | 7-9 | CASTLQRPGRREVVSQYF | 2-5 | moderate | CD4 | 9 |
| 20 | CAVRRGYGQNFVF | 26 | 7-9 | CASRRTSGRMRDNEQFF | 2-1 | strong | CD4 | 3 |
| 21 | CAVRPGFGNVLHC | 35 | 7-9 | CASSLARPFSGPQETQYF | 2-5 | moderate | CD4 | 3 |
| 21 | CAVRVGXGNVLHC | 35 | 7-9 | CASSRRRQGQETQYF | 2-5 | weak | DN | 1 |
| 21 | CAVKTGFGNVLHC | 35 | 7-9 | CASSAARLAGGSQFF | 2-1 | moderate | DN | 2 |
| 22 | CAVERLGAGGTSYGKLTf | 52 | 7-9 | CASSHRGRRRREYTEAFF | 1-1 | moderate | CD4 | 3 |
| 23 | CAAKGGGNKLTF | 10 | 7-9 | CASSIRGRVTEAFF | 1-1 | moderate | CD8 | 6 |
| 26-1 | CIVRVVYGKLVF | 47 | 7-9 | CASSPPPGTGVREQYF | 2-7 | weak | DN | 2 |
| 26-2 | CIXRPRAQKLVF | 54 | 7-9 | CASSRLQGLRSGKLFF | 2-7 | weak | CD4 | 6 |
| 30 | CGTRGGFGNVLHC | 35 | 7-9 | CASSTSAGRARSTEAFF | 1-1 | strong | DN | 4 |
| 35 | CAGPTRYNQGGKLIF | 23 | 7-9 | CASSLGRVGTSYNEQFF | 2-1 | weak | - | 8 |
| 38-1 | CAFRHQGAQKLVF | 54 | 7-9 | CASSLSRFGNQPHF | 1-5 | weak | CD4 | 9 |
| 21 | CAVGRSGSARQLTF | 22 | 11-2 | CASSLVKTGTSGIRGEQFF | 2-1 | weak | - | 8 |
| 8-2* | CVVSDNTGFQKLVF | 8 | 14 | CASSHRGYTRTSNTGELFF | 2-2 | weak | CD4 | 3 |
| 3 | CAVRQRGTGRRALTF | 5 | 18 | CASSPPTGDQPHF | 1-5 | weak | DN | 2 |
| 9-2 | CALSLNTGGFKTIF | 9 | 19 | CASSSVGSRTAPRKLFF | 1-4 | weak | - | 8 |
| 13-1 | CAASIKRMTGFQKLVF | 8 | 19 | CASRTRGRGDNQPHF | 1-5 | weak | CD4 | 3 |
| 20 | CAVRGQNFVF | 26 | 20-1 | CSARDLRTGGKYGYTF | 1-2 | strong | CD8 | 6 |
| 10 | CVVRFSGSARQLTF | 22 | 24-1 | CATSDLRSGQYF | 2-5 | weak | - | 8 |
| 12-3 | CAMSXLVRSNRKLIW | 37 | 28 | CASSPPRGDLSYEQYF | 2-7 | weak | CD4 | 3 |
| 17 | CATAGGTASKLTf | 44 | 28 | CASSSSYRYLLTDTQYF | 2-3 | weak | CD4 | 3 |
| 17 | CXTDGRGAQKLVF | 54 | 28 | CASSRPRYREKRTEAFF | 1-1 | moderate | DN | 3 |
| 36 | CAGSINTGTASKLTf | 44 | 28 | CASSSVRARSEKLFF | 1-4 | weak | - | 7 |
| 3 | CAVRDMIKAAGNKLTf | 17 | 30 | CAWSPIAGRRNEQFF | 2-1 | weak | - | 8 |

\**TRAV*8-2 or *TRAV*8-4\*\**TRBV*5-4 or *TRBV*5-8
X = amino acid not determined
Co-R = Co-receptor
- = co-receptor not determined

**Table S3.**
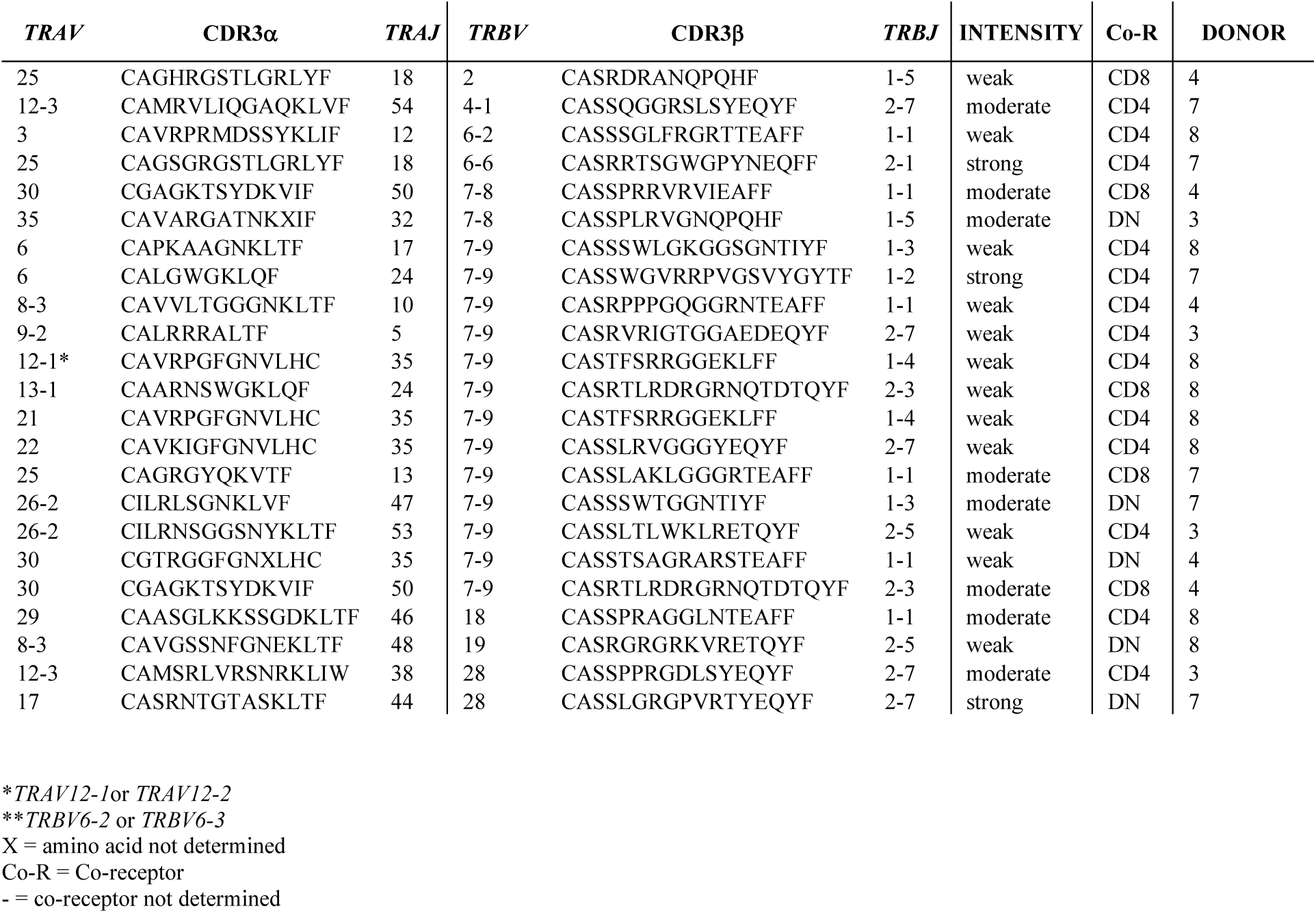
TCR sequences from CD1c-MPM tetramer^+^ αβ T cells.

| *TRAV* | CDR3α | *TRAJ* | *TRBV* | CDR3β | *TRBJ* | INTENSITY | Co-R | DONOR |
| --- | --- | --- | --- | --- | --- | --- | --- | --- |
| 25 | CAGHRGSTLGRLYF | 18 | 2 | CASRDRANQPQHF | 1-5 | weak | CD8 | 4 |
| 12-3 | CAMRVLIQGAQKLVF | 54 | 4-1 | CASSQGGRSLSYEQYF | 2-7 | moderate | CD4 | 7 |
| 3 | CAVRPRMDSSYKLIF | 12 | 6-2 | CASSSGLFRGRTTEAFF | 1-1 | weak | CD4 | 8 |
| 25 | CAGSGRGSTLGRLYF | 18 | 6-6 | CASRRTSGWGPYNEQFF | 2-1 | strong | CD4 | 7 |
| 30 | CGAGKTSYDKVIF | 50 | 7-8 | CASSPRRVRVIEAFF | 1-1 | moderate | CD8 | 4 |
| 35 | CAVARGATNKXIF | 32 | 7-8 | CASSPLRVGNQPQHF | 1-5 | moderate | DN | 3 |
| 6 | CAPKAAGNKLTF | 17 | 7-9 | CASSSWLGKGGSGNTIYF | 1-3 | weak | CD4 | 8 |
| 6 | CALGWGKLQF | 24 | 7-9 | CASSWGVRPVGSVYGYTF | 1-2 | strong | CD4 | 7 |
| 8-3 | CAVVLTGGGNKLTF | 10 | 7-9 | CASRPPPGQGGRNTEAFF | 1-1 | weak | CD4 | 4 |
| 9-2 | CALRRRALTF | 5 | 7-9 | CASRVRIGHTGGADEQYF | 2-7 | weak | CD4 | 3 |
| 12-1* | CAVRPGFGNVLHC | 35 | 7-9 | CASTFSRRGGEKLFF | 1-4 | weak | CD4 | 8 |
| 13-1 | CAARNSWGKLQF | 24 | 7-9 | CASRTLDRGRNQDTQYF | 2-3 | weak | CD8 | 8 |
| 21 | CAVRPGFGNVLHC | 35 | 7-9 | CASTFSRRGGEKLFF | 1-4 | weak | CD4 | 8 |
| 22 | CAVKIGFGNVLHC | 35 | 7-9 | CASSLRVGGGYEQYF | 2-7 | weak | CD4 | 8 |
| 25 | CAGRGYQKVTF | 13 | 7-9 | CASSLAKLGGGRTEAFF | 1-1 | moderate | CD8 | 7 |
| 26-2 | CILRLSGNKLVF | 47 | 7-9 | CASSWTGGNTIYF | 1-3 | moderate | DN | 7 |
| 26-2 | CILRNSGGSNYKLTF | 53 | 7-9 | CASSLTWLKLRETQYF | 2-5 | weak | CD4 | 3 |
| 30 | CGTRGGFGNXLHC | 35 | 7-9 | CASSTSAGRARSTEAFF | 1-1 | weak | DN | 4 |
| 30 | CGAGKTSYDKVIF | 50 | 7-9 | CASRTLDRGRNQDTQYF | 2-3 | moderate | CD8 | 4 |
| 29 | CAASGLKKSSGDKLTF | 46 | 18 | CASSPRAGGLNTEAFF | 1-1 | moderate | CD4 | 8 |
| 8-3 | CAVGSSNFGNEKLTF | 48 | 19 | CASRGGRKVRRETQYF | 2-5 | weak | DN | 8 |
| 12-3 | CAMSRLVRSNRKLIW | 38 | 28 | CASSPPRGDLSYEQYF | 2-7 | moderate | CD4 | 3 |
| 17 | CASRNTGTASKLTF | 44 | 28 | CASSLGRGPVRTYEQYF | 2-7 | strong | DN | 7 |

\**TRAV12-1* or *TRAV12-2*\*\**TRBV6-2* or *TRBV6-3*
X = amino acid not determined
Co-R = Co-receptor
- = co-receptor not determined

**Table S4.** MPM7-CD1c-PM CryoEM data collection and refinement statistics.

|  | MPM7 TCR-CD1c-PM ternary complex<br>EMDB-80696<br>PDB 26JD |
| --- | --- |
| **Data collection and processing** |  |
| Magnification | 105,000 |
| Voltage (kV) | 300 |
| Electron exposure (e-/Å<sup>2</sup>) | 60 |
| Defocus range (μm) | 0.5 - 1.5 |
| Pixel size (Å) | 0.82 |
| Symmetry imposed | C1 |
| Initial particle images (no.) | 514,401 |
| Final particle images (no.) | 130,855 |
| Map resolution (Å) | 2.90 |
| FSC threshold | 0.143 |
| Map resolution range (Å) | 2.90 - 3.20 |
| **Refinement** |  |
| Initial model used | PDB ID 6C15 (CD1c),<br>de novo (MPM7 TCR, Swiss-Model) |
| Model resolution (Å) | 3.2 |
| FSC threshold | 0.143 |
| Model resolution range (Å) | 2.8 - 3.1 |
| Map sharpening B factor (Å<sup>2</sup>) | 89.6 |
| Model composition |  |
| Non-hydrogen atoms | 6123 |
| Protein residues | 760 |
| Ligands | PM:1, GD3:1, NAG: 3 |
| B factors min/max/mean (Å<sup>2</sup>) |  |
| Protein | 7.64/121.17/46.72 |
| Ligand | 6.82/83.59/40.38 |
| R.m.s. deviations |  |
| Bond lengths (Å) | 0.004 |
| Bond angles (°) | 0.674 |
| Validation |  |
| MolProbity score | 2.63 |
| Clashscore | 12.66 |
| Poor rotamers (%) | 5.67 |
| Ramachandran plot |  |
| Favoured (%) | 93.03 |
| Allowed (%) | 6.69 |
| Disallowed (%) | 0.27 |

